# Dissecting chronic myeloid leukaemia overlapping transcriptome with TIF-Seq2

**DOI:** 10.1101/859488

**Authors:** Jingwen Wang, Bingnan Li, Sueli Marques, Lars M. Steinmetz, Wu Wei, Vicent Pelechano

## Abstract

Eukaryotic transcriptomes are complex involving thousands of overlapping transcripts. The interleaved nature of the transcriptome limits our ability to identify regulatory regions and, in some cases, can lead to misinterpretation of gene expression. To improve the understanding of the overlapping transcriptome, we have developed an optimized method, TIF-Seq2, able to sequence simultaneously the 5’ and 3’ ends of individual RNA molecules at single-nucleotide resolution. We investigated the transcriptome of a well characterized human cell line (K562) and identify thousands of unannotated transcript isoforms. By focusing on transcripts which are challenging to be investigated with RNA-seq, we accurately defined boundaries of lowly expressed unannotated and read-though transcripts putatively encoding fusion genes. We validated our results by targeted long-read sequencing and standard RNA-Seq for chronic myeloid leukaemia patient samples. Taking the advantage of TIF-Seq2, we explore transcription regulation among the overlapping units and investigate their crosstalk. We show that most overlapping upstream transcripts use poly(A) sites within the first 2 kb of the downstream transcription unit. Our work shows that, by paring the 5’ and 3’ end of each RNA, TIF-Seq2 can improve the annotation of complex genomes, facilitates accurate assignment of promoters to genes and easily identify transcriptionally fused genes.

**Key points:**

- Study of TSS-PAS co-occurrence allows dissecting complex overlapping transcription units.
- Partially overlapping transcription units in human commonly use PAS within the first 2Kb.
- TIF-Seq2 facilitates the identification of lowly expressed and transcriptionally fused genes.

## INTRODUCTION

Eukaryotic transcriptomes are complex, involving thousands of coding and non-coding RNA isoforms differing in transcription start sites (TSSs), poly(A) sites (PASs) and splicing. Overlapping isoforms can have divergent functional consequences, changing the encoded protein (1, 2) or affecting mRNA post-transcriptional life (e.g. translation, localization and stability) (3). However, the interleaved nature of the transcriptome convolutes its study and limits the accurate identification and quantification of alternative isoforms (4, 5). This can lead to incomplete or inaccurate annotations, which cause misinterpretation of gene expression data (6) and limits our ability to link regulatory regions with genes (7) (and thus to genetically manipulate them). Particularly challenging is the correct identification of transcription boundaries of overlapping isoforms (8), even if we know that alternative TSS and PAS drive most isoforms variations across human tissues (9). Standard RNA-Seq can identify transcribed regions and splicing events, however, it cannot distinguish RNA fragments originating from alternative overlapping features. The overlapping nature of the transcriptome also limits our ability to dissect the molecular mechanism underlaying the regulatory crosstalk across adjacent transcriptional units (10, 11). For example, RNA-Seq cannot distinguish if a specific mRNA fragment in the 5’region of a gene will reach the canonical poly(A) site or originate from an overlapping transcriptional unit that terminates prematurely. Single-end approaches such as CAGE or poly(A) site sequencing have been key to define the boundaries of the transcriptomes (12, 13). However, those approaches cannot study the combination between TSS and PAS. Long-read sequencing technologies promise to reveal transcription complexity on the genome-wide scale (14). Nevertheless, the high cost it entails combined with low throughput and limited resolution in the 5’ and 3’ transcript regions are still major limitations (15).

To bridge the gap between short-read and long-read technologies, and to improve our ability to study the regulatory crosstalk between same strand overlapping transcriptional units, we have developed an optimized **T**ranscript **I**so**f**orm **Seq**uencing (TIF-Seq2) that is especially well-suited for the interrogation of complex transcriptomes. TIF-Seq2 allows to sequence simultaneously the 5’ and 3’ end of individual RNA molecules at single-nucleotide resolution. To demonstrate its utility, we dissected the overlapping transcriptome of a chronic myeloid leukaemia (CML) cell line (*i.e*. K562) in response to Imatinib treatment. We identify thousands of known and unannotated transcript isoforms, accurately define the boundaries of lowly expressed intergenic transcripts and validate them using alternative short-read and targeted long-read sequencing approaches. We focus on overlapping transcriptional units that are particularly challenging to investigate with RNA-Seq, and show the common existence of short overlapping upstream transcripts that may lead to misinterpretation of RNA-Seq and CAGE gene expression data. We also show the existence of more complex overlapping and read-through transcripts. Finally, we used the obtained information to improve the detection and analysis of complex overlapping transcripts in clinical RNA-Seq datasets and show evidences of transcriptional events involving gene-promoter rewiring and potentially leading to the generation of transcriptionally fused proteins.

## MATERIALS AND METHODS

### Cell Culture

The human erythroleukemia cell line K562 was obtained from ATCC(ATCC^®^CCL-243™). Cells were cultured in RPMI 1640 medium supplemented with 10% FBS, 2 mM L-glutamine, 1% pen/strep (Life Technologies) at 37 °C with 5% CO2 in a humidified atmosphere. Cells (3 X 10^5^ cells/mL) were exposed to 0.2 to 5μM (0.2, 0.5, 1, 1.7 and 5μM) imatinib for 8h, 24 h and 48h. Aliquots were taken at each time point for assessment of cell viability via Trypan blue staining by EVE™ automated cell counter. Two biological replicates of K562 cells treated with 1μM imatinib for 24h and corresponding DMSO control were used for TIF’Seq2 library preparation.

### TIF-Seq2 Library preparation

In brief, capped and polyadenylated mRNA was used as template to generate full-length cDNA. We circularized the cDNA, remove non circularized molecules and fragmented circularized molecules using sonication. Streptavidin magnetic beads were used to purify the fragments spanning the 5’ and 3’ end of cDNA and then were used for Illumina library preparation (Additional file 1: Figure S1). In detail, 2.5 μg total RNA was treated with Turbo DNase (0.12U/μl) (Fisher Scientific) for 20min at 37°C to prevent genomic DNA contamination. After inactivation of DNase, input RNA was dephosphorylated by incubating with Calf Intestinal Alkaline Phosphatase (0.3U/μl) (CIP)(NEB) at 37°C for 30 min. Two rounds of phenol-chloroform extraction followed by ethanol precipitation was performed to remove CIP. After dephosphorylation, input RNA was decapped by incubating with Cap-Clip Acid Pyrophosphatase (0.125U/μl) (CellScript) at 37°C for 60min. After phenol-chloroform purification and ethanol precipitation, RNA was ligated overnight at 16°C at 5’ with DNA/RNA chimeric oligonucleotide adaptor (TCAGACGTGTGCTCTTCCGATCTrNrNrWrNrNrWrNrN, TIF2-RNA in Supplementary Table S1 using T4 RNA ligase (NEB) in the presence of 10% dimethylsulphoxide (DMSO), RiboLock RNase Inhibitor (Thermo Fisher Scientific EO0382) and 1mM ATP. The chimeric TIF2-RNA adaptor introduced a common anchor sequence for forward primer of subsequent PCR amplification and an 8-mer unique molecular identifier (UMI). Ligated RNA was purified with 1.8:1 volumetric ratio (1.8X) RNA clean XP beads according to the manufacturer instruction and then used as the template for reverse transcription (RT). Ligated mRNAs were reverse transcribed using barcoded oligo-dT primers (i.e., TAGTTCAGTCTTCAGTACCTCGTGCGGCCGCXXXXXXACACTCTTTCCCTACACGACGCTCTTCCGATCTIIIIIIIIIIIIITTVN; where X refers to the specific barcode, TIF2-RT in Supplementary Table S1) which introduced Illumina sequencing primer 1, a 3’ index, a Not1 endonuclease digestion site and a 3’ common sequence for the subsequent PCR reaction. Specifically, mRNAs were mixed with the corresponding TIF2-RT oligo and dNTPs, denatured at 65°C for 5min and then put on ice. The sample was mixed with 5x First strand buffer, Trehalose (1.57M) and RiboLock RNase inhibitor and incubated first at 42°C for 2 min. Finally, 2 μl of SuperScript™ III reverse transcriptase (Thermo Fisher Scientific) was added to each reaction and incubated at 42°C for 50 min, at 50°C for 30 min, at 55°C for 30 min and inactivated at 70°C for 15 min. Used RNA template was removed by incubating with 0.5ul RNase H(5U/μl) and 0.5μl RNase cocktail (Ambion) at 37°C for 30min. First-strand cDNA was purified with 2X Ampure XP beads according to manufacturer instruction. To avoid saturating the reaction, only half of the obtained cDNA was used for the following PCR, and the rest was stored as a backup. Used cDNA template was further split into two PCR reactions with Terra PCR Direct Polymerase (Takara) with the following program 98°C for 2min, then 16 cycles of 98°C for 20s, 60°C for 30s, 68°C for 5min (+10s/cycle) and finally 72°C for 5min. The PCR above used as primers TIF2-Rv: TAGTTCAGTCTTCAGTACCTCGT and TIF2-Fw: TATAGCGGCCGCXXXXXXGTGAC[BtndT]GGAGTTCAGACGT GTGCTCTTCCGATC (where X refers to different barcodes). PCR products were purified with 1X Ampure XP beads according to manufacturer instruction and then quantified with Qubit dsDNA HS assay. PCR products from different samples were then pooled together with equal mass. By pooling full length cDNA containing sample specific barcodes, we were able to detect and estimate the percentage of intermolecular circularization events (i.e. chimeras connecting barcodes originating from different samples). This information was used to select the optimal concentration favouring intramolecular ligation (see below). Pooled PCR products were subjected to 1U/μl Not1 HF (New England BioLabs) endonuclease digestion at 37°C for one hour. After inactivation of Not1 at 65°C for 20 min, samples were purified with 1.8x Ampure XP beads according to manufacturer instruction.

To favour intramolecular ligation, PCR products with sticky ends were highly diluted to a final concentration less than 1ng/μl and ligated with a high concentration (66.68U/uL) of T4 DNA ligase (New England BioLabs) at 16°C for at least 16 hours. To remove the unligated linear PCR products, we added 0.5 μl plasmid-safe for every 100 μl total volume and incubated at 37°C for 1h 1h in the presence of 1uL ATP 100mM. After inactivation at 70°C for 30min, the self-circularized cDNA was purified with phenol-chloroform and ethanol precipitation. Circularized cDNA was fragmented by sonication (Covaris ME220; Covaris, Inc) (Duration 240s, peak power 30, duty factor 10, 200 cycles/burst, Avg Power(W) 3. The fragments were then purified with 1X Ampure XP beads according to manufacturer instruction. Biotin labelled fragments which contained the 5’ and 3’ connecting region was enriched using Dynabeads M280 streptavidin (Invitrogen) and incubating at room temperature for 30 min. Captured fragments were subjected to end repair with End Repair Enzyme Mix (New England Biolabs) and dA tailing with Klenow Fragment exo 5U/uL (New England Biolab). To add the required Illumina grafting sequences, each sample (20μl of resuspended beads) was incubated at room temperature for one hour with a mixture of 10 μl 5X Quick ligation buffer (New England BioLabs), 16μl nuclease-free water (Ambion), 3 μl T4 DNA ligase(2000U/μl) and 1 μl 1μM duplex adaptors. Duplex adaptors were generated by annealing TIF2-forkFw (5’-AATGATACGGCGACCACCGAGATCTACACACACCTGCCGGTCACC*T-3) and TIF2-forkRv (5’-phos-GGTGACCGGCAGGTGTATCTCGTATGCCGTCTTCTGCTTG-3’) at 15μM and diluted to 1μM working solution freshly when used each time. After cleaning the beads, the beads resuspended in 20μL EB buffer were used for PCR amplification using 25μl Phusion High-fidelity MasterMix (2X) (New England BioLabs), 4 μl nuclease-free water and 0.5 μl PCRgraftP5 primer (5uM, 5’-AATGATACGGCGACCACCGAGATCTACAC-3’) and 0.5 μl PCRgraftP7 primer (5uM, 5’-CAAGCAGAAGACGGCATACGAGAT-3’). Samples were subjected to PCR amplification with the following program: 98°C for 30s, then 18 cycles of [98°C for 20s, 65°C for 30s, 72°C for 30s] and finally 72°C for 5min. DNA library then went through two step beads-based size selection (first step 0.35X, second step take the supernatant from first step and add extra 0.45X) with expected size distribution in the range of 300bp-1000bp. Purified DNA library was adjusted to 4nM, then denatured, diluted according to Illumina instruction and sequenced on Nextseq 500 platform.

We used four custom sequencing oligos as follows: SeqR1+15T (5’-ACACTCTTTCCCTACACGACGCTCTTCCGATCTTTTTTTTTTTTT TTT-3’), SeqINDX1 (5’-GATCGGAAGAGCACACGTCTGAACTCCAGTCAC-3’), SeqINDX2 (5’-GATCGGAAGAGCGTCGTGTAGGGAAAGAGTGT-3’) and SeqR2 (5’-GTGACTGGAGTTCAGACGTGTGCTCTTCCGATCT-3’). See Additional file 1: Figure S2 for details regarding the sequencing oligonucleotide annealing. Two mixtures of sequencing oligos were prepared as follows: mixture 1 contained SeqR1+15T(0.3μM) and SeqR2(0.3μM), mixture 2 composed of SeqINDX1 primer (0.3μM) and SeqINDX2 primer (0.3μM). Mixture 1 was loaded into both positions (#7 and #8 on reagent cartridge in a NextSeq 500 instrument) for custom read1 primer and for custom read2 primer. Mixture2 was loaded into the position for custom index primer (#9 on reagent cartridge). Sequencing was carried out in an Illumina NextSeq 500 instrument with stand-alone configuration and custom sequencing oligos. Paired end sequencing read lengths were set as read1 76bp, read2 76bp, index1 6bp and index2 6bp.

### 3’T-fill library preparation

We performed 3’T-fill as previously described (16). In brief, 10μg DNA-free total RNA (16μL) was fragmented by adding 4μl fragmentation buffer(5X)(200mM Tris-acetate, pH 8.1, 500mM Potassium Acetate, 150mM Magnesium acetate) and incubated at 80°C for 5min. Fragmented RNA was purified with 1.5X Ampure XP beads according to manufacturer instruction and used as the template for reverse transcription(RT). Sample were mixed with biotinylated dT primer (P5_dT16VN 5’-[Btn]AATGATACGGCGACCACCGAGATCTACACTCTTTCCCTA CACGACGCTCTTCCGATCTTTTTTTTTTTTTTTTVN-3’) (where V refers to A, C or G) and denatured at 65°C for 5min. Denatured RNA was then mixed with 4μl 5X first strand buffer, 2μl DTT(0.1M) and 4μl freshly prepared actinomycin D(0.1μg/ul). After incubation at 42°C for 2min, 0.5μl SuperScript™ II (Invitrogen) was added, incubated at 42°C for 50 min and finally 72°C for 15min. cDNA was purified with 1.5X Ampure XP beads according to manufacturer instruction. For second strand synthesis, cDNA was mixed with of 0.5 μl RNase H (5U/μl) (New England BioLabs) and 2 μl DNA Polymerase I (10U/μl) (Thermo Scientific) and incubated at 16°C for 2.5 hours. cDNA was purified with 0.9X Ampure XP beads according to manufacturer instruction. 20 μl biotin labelled cDNA was mixed with 20μl Dynabeads M280 streptavidin beads (Invitrogen) at room temperature for 15 min. Following end repair and dA tailing, each sample(8 μl) was mixed with 12.5 μl 2X Quick Ligation buffer (New England BioLabs), 2.5μL T4 DNA ligase (2000U/μL, New England BioLabs) 2μl of anneal duplexed adaptors at 2.5 μM(5’-GTGACTGGAGTTCAGACGTGTGCTCTTCCGATC*T-3’, 5’-Phos-GATCGGAAGAGCACACGTCTGAACTCCAGTCAC[AmC7]-3’) and incubated for at 20°C for 20min. Adaptor ligated libraries were washed and amplified by PCR by adding 25 μl 2X Fusion High Fidelity Master Mix, 0.5 μl PE1 primer (10μM, 5’-AATGATACGGCGACCACCGAGATCTACACTCTTTCCCTACAC GACGCTCTTCCGATC*T-3’)and 0.5μl PE2_MPX primers (10μM, 5’-CAAGCAGAAGACGGCATACGAGATXXXXXXGTGACTGGAGT TCAGACGTGTGCTCTTCCGATC*T-3’; where X refer to the specific barcode). Samples were subjected to the following program: 98°C for 30 seconds, 18 cycles of 98°C for 10s, 65°C for 10s, 72°C for 10s and finally 72°C for 5min. Library was purified with 1.8X Ampure XP beads according to manufacturer instruction.

### PacBio long reads sequencing

To validate the existence and structure of unannotated transcript isoforms, we used the TIF-Seq2 derived information to perform targeted amplification followed by Pacific Biosciences (Pac Bio) sequencing. To decrease the costs associated to library preparation, we performed a pooled reverse transcription with a mix of gene specific primers and then split the obtained cDNAs for individual PCR reactions. We used as input 2 biological replicates of K562 exposed or not to imatinib (4 samples, as in the TIF-Seq2). Each sample was individually labelled during the PCR reaction (see below). We pooled equal amount of all isoform-specific RT primers (Supplementary Table S1) and used total RNA as a template for reverse transcription (RT). Isoform-specific RT primers (PB_RT) were designed with a universal primer sequence (5’-GTGACTGGAGTTCAGACGTGT), plus 8 random nucleotides as unique molecular identifier and a gene specific sequence. RNA was mixed with primer and dNTPs, denatured at 65°C for 5min and transferred to ice. For the reverse transcription we added first strand buffer, Trehalose (1.57M), RiboLock RNase Inhibitor and the mix was incubated at 42°C for 2min, and then 2 μl of SuperScript II reverse transcriptase (ThermoFisher Scientific) was added to each reaction. We incubated each reverse transcription reaction at 42°C for 50min, 50°C for 30 min, 55°C for 30min and inactivated at 70°C for 15min. Obtained cDNA was distributed in 96-well plates containing in each well an isoform-specific forward primer (PB_FW, composed of a common sequence 5’-ACACTCTTTCCCTACACGAC and gene specific sequence) and a common reverse primer containing 8 mer sample identifier (FigS13). cDNA was PCR amplified using the Phusion High Fidelity Master Mix with the following program: denaturation at 98°C for 2min, then 98°C 20s, 60°C 30s, 68°C 5min (+10s/cycle) for 35 cycles and final extension was performed at 72°C for 5min. Amplified products were analysed by gel electrophoresis, purified by 1.8X Ampure XP beads, and pooled at similar concentration for sequencing. Pooled PCR products were used for library generation using SMRTbell™ Template Prep Kit 1.0-SPv3 and sequenced on PacBio Sequel system.

### TIF-Seq and 3’T-fill sequencing processing and alignment

We employed bcf2fastq (v2.20.0) for converting raw image to sequence information (FASTQ files) and demultiplex, allowing two mismatches in index 1 and one mismatch in index 2. For TIF-Seq2 data, we collapsed all 5’-end or 3’-end sequence reads in each sample according to the indexes. Cutadapt (17) (v1.16) was utilized to trim TIF-seq2 sequencing primer (-a AGGTGACCGGCAGGTGT) and Illumina TruSeq adapter (-a AGATCGGAAG). After extracting 8 bp of unique molecular identifiers (UMIs) with UMI-tools (18) (v0.5.4) from the 5’ ends and removing extra A stretches in the 3’ ends caused by poly(A) slippage during PCR amplification, we kept reads over 20 bp for alignment. We used STAR (19) (v2.5.3a) for aligning 5’-end reads and 3’-end reads separately to the human reference genome hg38, supplying Gencode v27 transcripts as splicing junction annotation. Alignment setting was adjusted as below, --alignIntronMax 200000 --alignEndsType Extend5pOfRead1 --alignSJoverhangMin 10. We then linked paired-end reads and kept the uniquely mapped pairs that are on the same chromosome (Additional file 1: Figure. S4). Furthermore, a customised script adapted from UMI-tools was employed to remove PCR duplicates from the leftover reads, allowing 1 bp mismatch in the UMIs and 1 bp shifting in the transcription start sites (https://github.com/jingwen/TIFseq2/blob/master/dedup.py).

For 3’T-fill sequencing data, we trimmed Illumina TruSeq adapter (-a AGATCGGAAG) and extra A stretches in the 3’ ends with Cutadapt v1.16. Reads over 20 bp are aligned to hg38 by using STAR v2.5.3a in paired-end mode, supplying Gencode v27 transcripts as splicing junction annotation with adjusted setting, --alignSJDBoverhangMin 1 --alignIntronMax 1000000 --alignMatesGapMax 1000000 --alignEndsType Extend5pOfReads12. Only uniquely mapped reads were kept for downstream analysis.

In order to evaluate the improvement of TIF-Seq2, we compared our data from the current study to a previous study on human HeLa cells using TIF-Seq1 (20) (GEO: GSE75183). We aligned TIF-Seq1 reads to the human reference genome hg38 using STAR with the same parameter setting as how we analysed TIF-Seq2 data.

### Transcription boundary determination

A customised python script was employed to extract both boundaries of TIF-Seq2 read pairs and 3’ tags of 3’T-fill sequencing data, collapse the boundary tags and calculate the coverage (https://github.com/jingwen/TIFseq2/blob/master/boundary.py). In order to filter out false positive poly(A) sites caused by internal priming, we exclude the 3’ end tags with at least 7 As in the downstream 10 nt sequences. Then we employed CAGEr (21) to define the cluster of transcripts 5’- or 3’-end tags of TIF-Seq2 respectively (Additional file 1: Figure S4b). Transcription boundary tags were normalized to match a power-law distribution. Low-coverage tags supported by less than 1 normalized counts in more than 1 sample were excluded before clustering. The boundary tags within 10 bp window were spatially clustered together. Clusters with only one boundary tag are kept if the normalized counts are above 1. The tag clusters were further form into non-overlapping consensus clusters across all samples if they are within 10 bp apart. The same strategy was applied for identifying consensus PAS cluster from 3’T-fill sequencing data.

### TIF definition, annotation and quantification

We then link the 5’-end TSS and 3’-end PAS clusters according to the supporting read pairs from TIF-Seq2 (Additional file 1: Figure S4b). We filtered out pairs with extremely long (> 2Mb) and extremely short (< 300 bp) mate-pair distance. To keep a conservative estimate of unannotated transcript isoforms identified, we excluded pairs mapping to different chromosomes. Transcript isoform boundaries (TIFs) were defined as connection between TSS clusters and PAS clusters supported by at least 4 read pairs connecting them across all samples (unique molecular events). The TIFs were further assigned to Gencode v28 annotation features based their relative distance to the annotated transcripts (Additional file 1: Figure S4c). TSS distance (d1) and PAS distances (d2) are calculated between a TIF and its overlapping annotated transcripts. The TIF is assigned to the transcript with the least sum of d1 and d2 among all overlapping transcripts, further assigned to the gene that harbour the transcript. According to the relative position to their assigned transcripts, the TIFs were classified as 1) annotated transcripts, if both TSS and PAS are within 200 bp away of annotated transcripts boundaries; 2) transcripts with new TSS; 3) transcripts with new PAS; 4) transcripts with new boundaries, if both TSS and PAS not annotated and 5) intergenic TIFs (Additional file 1: Figure S9). We measured TIF-Seq2 expression in K562 cells as count of read pairs that link TIF boundaries. We employed DEseq2 (22) for normalization and differential expression analysis (before and after drug treatment) with default setting.

### Long-read sequencing data analysis

We trimmed the PCR primers from both ends of the PacBio highly accurate consensus sequences using Cutadapt (v1.16) and extract UMIs with UMI-tools (v1.0.0), keeping the reads with at least 200 bp in length. Then the reads were aligned to human reference genome using minimap2 (23) (v2.16) with the following setting (-ax splice -uf -C5 -O6,24 -B4). We used UMI-tools for removing the PCR duplicates with adjusted setting as --method cluster --spliced-is-unique. We further employed BEDTools (24) (v2.27.1) bamToBed function to convert the alignment reads into BED format and a customised script to convert BED format to GFF format.

### Independent RNA-Seq validation

RNA-Seq data of K562 cells before and after Imatinib treatment (n=4) from study Gallipoli et al. (25) were download from GEO depositories (GSE105161). RNA-Seq data of 21 paired CML patient (26) before and after Imatinib treatment from were downloaded from EGA archive (EGAD00001004179). We employed STAR (v2.5.3a) to align paired-end reads to human reference genome hg38, adding long-read sequencing validated transcripts into splicing junction annotation, with adjusted setting (--alignSJDBoverhangMin 1 --alignIntronMax 200000 --alignEndsType Extend5pOfReads12). Polyribosome profiling data of K562 cells (27) (n=3) was downloaded from GEO (GSE93210). We used HISAT2 (28) (v2.1.0) for aligning the reads to human reference genome hg38 and long-read sequencing validated transcripts as splicing junction annotation, meanwhile adjusting maximum intron length to 200kb.

Gene expression in K562 cells (25) and CML patients (26) from standard RNA-Seq was quantified using featureCounts (29) according to Gencode v28 transcript annotation, TIF-Seq2 transcription boundaries and long-read sequencing validated transcript model. We employed DEseq2 (22) for normalization and differential expression analysis (before and after drug treatment) with default setting for K562 cells. For CML patient RNA-Seq data, we set up multiple factors (paired samples, drug treatment and phenotype) in DESeq2 for differential expression test.

## RESULTS

### TIF-Seq2 delineates isoforms in complex human transcriptome

We and others have previously developed approaches able to link the 5’ and 3’ regions of individual transcripts (30–32). Our work in *S. cerevisiae* demonstrated the existence of a complex overlapping transcriptome even in a simple eukaryote with limited splicing (30). However, the applicability of our original protocol (33) to the study of complex transcriptomes was limited due to the variability in length of the mappable 5’ and 3’ tags and its modest throughput. To increase the length of the boundary tags, we designed a new sequencing strategy decoupling the region required for bridge amplification from the Illumina sequencing primers (Figure. 1A; Supplementary Figure S1-2). This decreases the need to perform stringent library size-selection, allows sequencing from the exact 5’ and 3’ ends of each mRNA molecule and generates longer reads suitable for the study of complex genomes (see Methods). In addition, we performed extensive enzymatic optimizations to maximize the length and complexity of the full-length cDNA libraries, as well as introduced early sample pooling, unique molecular identifiers (UMI) and barcodes to control for the formation of chimeras (Supplementary Figure S1).

**Figure 1.**
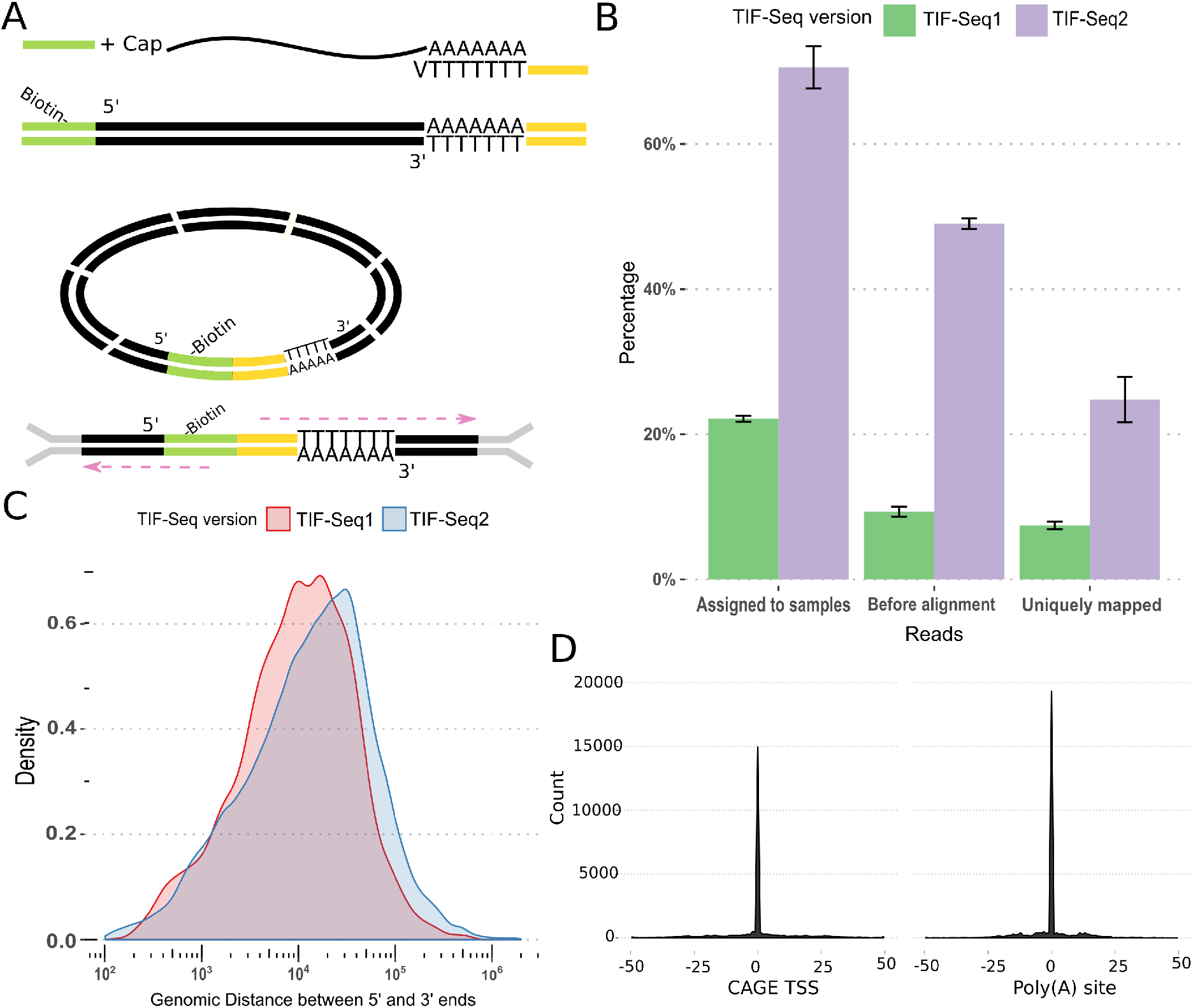
Genome-wide measurement of transcript isoforms with TIF-Seq2. (A) TIF-Seq2 protocol; (B) Informative reads fetched from TIF-Seq1 (n=3) and TIF-Seq2 (n=4). TIF-Seq2 can fetch more useful reads that are assigned to samples, reads passed quality control for alignment and the uniquely mapped reads; (C) Genomic distance between 5’ and 3’ ends captured by TIF-Seq1 and TIF-Seq2. The enzymatic optimization of TIF-Seq2 can improve the lengths of RNA molecules (average distance: 20 kb in TIF-Seq1 and 35 kb in TIF-Seq2); (D) Transcript boundaries agree with the transcription start sites (TSSs) defined by CAGE (12) and the poly(A) sites defined by 3’ sequencing (13).

To demonstrate its utility, we investigated the transcriptome of a well-characterized CML cell line (K562) in response to the tyrosine kinase inhibitor Imatinib (Supplementary Figure S3). After quality control and PCR deduplication (see Methods and Supplementary Figure S4A), we obtained over 14 million pair tags uniquely mapped to the human genome at single-nucleotide resolution (Supplementary Table S2-3). Compared with TIF-Seq1 (20), all these modifications improved the number of informative reads, genomic distances of transcription boundaries (Figure 1B-C), and thus allowed the application of TIF-Seq2 to complex genomes. We clustered adjacent transcription start sites (TSSs) and polyadenylation sites (PASs) and obtained 32,631 TSS and 31,187 PAS clusters (see Methods). Identified clusters are narrow, and 80% of them are smaller than 15 nt (TSS) and 10 nt (PAS) (Supplementary Figure S5A and B). To validate the accuracy of TIF-seq2, we performed 3’T-fill (16) with the same samples and compared transcript boundaries with independent published datasets for TSS (CAGE (12)) and PAS measurements (13). In both cases, those analysis demonstrates our accurate detection of transcript boundaries (Fig. 1D and Supplementary Figure S6). Among high-confidence (normalised counts > 10) TIF boundaries, 85% of TSS and 92% of PAS clusters are located within 100 bp next to the TSSs and PASs from the published datasets (12, 13). The main advantage of our approach is that it allows to link the identified TSS and PAS clusters (Supplementary Figure S4). Doing so, we identified 49,847 unique combinations of TIF-seq2 linked TSS-PAS clusters (referred here as **TIF**s, *boundary* transcript isoforms) supported by at least 4 independent molecular events across all samples. TIF-Seq2-derived transcript boundaries are in good agreement with the curated Gencode v28 annotation (Supplementary Figure S7). At our current sequencing coverage, TIFs overlap 9006 annotated genes (80% covered by more than one TIF) (Supplementary Figure S8). In spite of the good agreement (Supplementary Figure S7), 60% of the TIFs support non-annotated isoforms with alternative TSSs, PASs, or both (Supplementary Figure S9). Current genome annotation is based mainly on classical cloning strategies, RNA-Seq and CAGE (5, 12), which limit its ability to distinguish between overlapping isoforms harboured in the same gene locus. As TIF-Seq2 is able to reveal complex overlapping transcript structures, we decided to focus on those that are often missed by traditional approaches.

### TIF-Seq2 improves transcript boundary delineation in clinical RNA-Seq data

Lowly expressed transcripts are in general challenging to detect. Long-read approaches lack the throughput, and even when combined with capture-based enrichment, they require a prior definition of the regions of interest (34). On the other hand, short-read RNA-Seq approaches with much higher throughput distribute their sequencing power along the whole transcribed region, and need to be linked with independent TSS or PAS dataset to accurately infer their putative boundaries (8). On the contrary, TIF-seq2 focus its sequencing power on the transcript boundaries (TSS and PAS), allowing to link them confidently even with relatively low coverage. This simplifies the annotation of lowly expressed transcripts and makes it easy to distinguish them from background noise. Using this approach, we identified 1034 TIFs in 426 unannotated non-overlapping transcribed regions, defined as unannotated genes (Supplementary Table S4). Although the main application of TIF-Seq2 is the dissection of the overlapping transcription organization, it also allows the quantification of differential expression of the same isoforms across samples. To test that, we treated the K562 cells with 1μM imatinib for 24h. Doing so, we identified 879 TIFs that are up/down-regulated (Supplementary Figure S10 and Supplementary Table S5). And among those, 60 TIFs in 36 unannotated non-overlapping transcripts were differential expressed (in agreement with our estimation by 3’T-fill, Supplementary Figure S11 and Supplementary Table S6). We found evidence for their transcription and expression pattern using independent RNA-Seq datasets (25) (Supplementary Table S4). About 24% of unannotated transcripts are significantly regulated (adjusted p-value < 0.05) after Imatinib treatment. In order to validate the identified isoforms and determine their internal structure, we used the TIF-Seq2-derived boundaries of 28 differentially expressed unannotated genes for designing amplicon-based enrichment of fulllength isoforms followed by long-read sequencing (Supplementary Figure S12). We confirmed the existence of 25 candidates and predicted the existence of short (29-179 aa) open reading frames in 17 cases (Figure 2A and Supplementary File 1), suggesting their coding potential.

**Figure 2.**
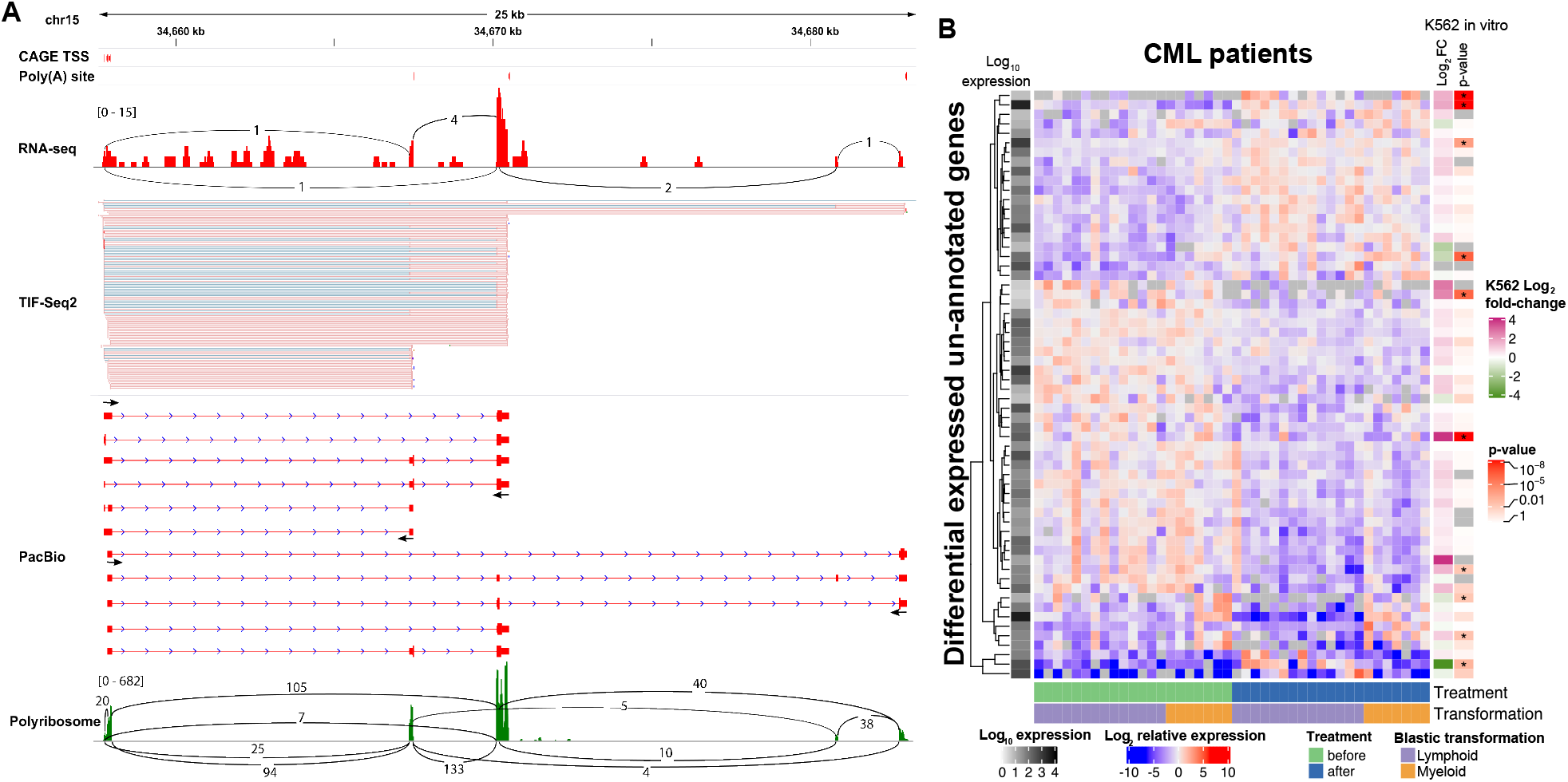
Unannotated lowly expressed intergenic transcripts can be detected in CML patients. (A) TIF-Seq2 can identify lowly expressed transcripts, for example, an unannotated gene on chr15: 34657737-34683026. TIF-Seq2 read pairs are labelled as pink lines (the forward strand). The fine blue lines in the TIF-Seq2 track represent splicing junctions in the first or last exons. CAGE TSS and poly(A) sites from public repositories (12, 13) and an independent K562 RNA-Seq experiment (25) validates the expression of this gene. Primer designed for target long-read Pac Bio validation are marked by arrows. Polyribosome-associated RNA-Seq (27) track is in the bottom, with expression marked in green and splicing junction in black lines, suggests its coding potential. (B) expression of unannotated transcription features in CML patient (26). Each row represents a differentially expressed gene (adjusted p-value < 0.005) in CML patient. Each column represents a sample before or after drug treatment. Patients were classified as lymphoid blast crisis (LBC) or myeloid blast crisis (MBC) according to the types of their blastic transformations. Each cell represents log_2_-scale ratio of gene expression in each sample to the average expression of the gene across all samples. The average expression (in log_10_ scale) of genes is depicted on the left. On the right side, we present the log_2_-scale fold change of those genes in an independent *in vitro* experiment exposing K562 cells to imatinib (25) and their significant levels of differential expression after treatment. Genes in K562 cells with adjusted p-value < 0.01 are labelled in asterisks.

After annotating those intergenic transcripts in K562, we investigated their potential relevance in chronic myeloid leukaemia (CML) patients. We used the TIF-Seq2 derived annotation to re-analyse RNA-Seq data from a cohort of CML patients responded to first-line TKI imatinib therapy, and focus on those who failed to achieve durable major molecular response and eventually developed blast crisis (26). We confirm that 365 unannotated genes identified in K562 are also expressed in at least three CML patients with minimum one count per million reads. 59 of them responded to Imatinib treatment, and 10 are differentially regulated between patient groups that developed myeloid blast crisis (MBC) or lymphoid blast crisis (LBC) (Figure 2B, adjusted p-value < 0.005). Interestingly transcripts downregulated by Imatinib treatment in K562 are upregulated in patients that stopped responding to treatment and underwent blast crisis. As an example, a K562-specific transcript on chr6:15555763-15559977 (putatively encoding the HTH domain of the Mos1 transposase) was significantly upregulated in patients who developed LBC after therapy, while it is significantly downregulated in K562 cell line after Imatinib treatment (Supplementary Figure S13A-C). Independently of the potential relevance of the newly identified transcripts for disease progression, it is clear that they are present but excluded from most analysis due to incomplete transcriptome annotation and difficulty of its analysis.

### Disentangling overlapping transcription units allows unequivocal assignment of promoter proximal poly(A) sites

After showing the ability of TIF-Seq2 to better identify the boundaries of lowly expressed transcripts, we focused on the analysis of overlapping transcription units. We first asked how common was the overlap between neighbouring same-strand transcription units and their degree of overlap. 40% of overlapping TIF pairs present a high degree (over 90%) of overlap (top right corner in Figure 3A). This represents alternative transcription isoforms in same genes with slightly different TSSs or PASs in the first or last exons. About 50% of the TIF pairs represent the overlap of a longer isoform and its truncated short isoform, which starts from an intronic TSS or ends at an intronic PAS (top or right side in Fig 3A respectively). The rest of the overlapping TIF pairs correspond to tandem short isoforms originating from either the same gene loci or tandem transcripts originating from upstream genes. As partial overlapping between transcriptional units offer clear opportunities for regulatory crosstalk among adjacent genes (11, 35) we decided to focus on those.

**Figure 3.**
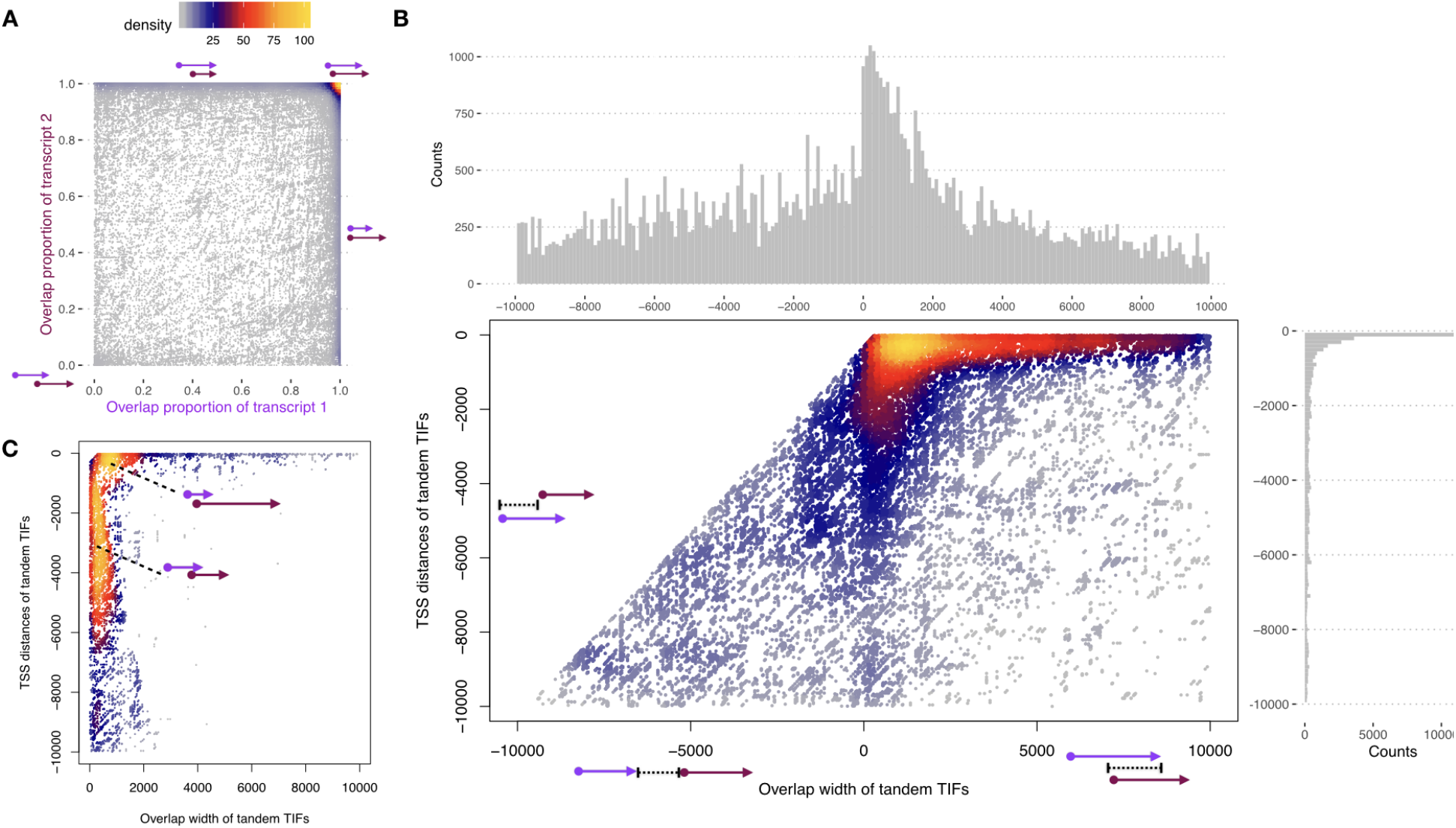
TIF-Seq2 reveals overlapping transcription units. (A) Pairwise analysis of the genomic overlap between TIFs. TIFs from the same gene loci present a high degree of overlap (over 90%, top-right corner), while TIFs from neighbouring tandem genes overlap at low degree (bottom-left corner). (B) Tandem TIFs captured by TIF-Seq2 reveal the path of RNA polymerase II (Pol II). Each dot in the heat scatter plot represents a pair of tandem TIFs within 10 kb. X axis shows overlap width between two tandem TIFs. In the case of no overlap, the values in X axis is the distance between two TIFs in the tandem TIF pairs. Y axis represents the distance of upstream TSS to the downstream TSS of tandem TIFs. Tandem TIFs typically overlap by less than 2 Kb (histogram on the top). The majority of the first TIFs originated from within 1 Kb upstream of the second TIFs. (C) Low degree overlapping tandem TIFs. Dots, x axis and y axis are as in Figure 3B.

First, we focus on overlapping TIFs originating within 10 Kb of each other. We can observe that upstream overlapping transcriptional units tend to use a poly(A) site located within the first 2 Kb of the downstream transcriptional unit (x-axis in Figure 3B). And among the overlapping tandem TIFs originating in a 10 Kb window from the downstream transcription unit, 60% of them arise within 1 Kb upstream of the second TIFs (y-axis in Figure 3B). This suggest that in those regions, even with constitutive nuclear RNA degradation machinery, coexist RNA polymerases originating from upstream promoters will terminate shortly and RNA polymerases will proceed until the canonical poly(A) site. When investigating the case of low-degree tandem overlapping TIFs (< 20% overlap in either TIF), we discovered two typical types of overlap (Figure 3C). One pattern (23%) is an upstream short TIF (< 2 Kb) originating from 1 Kb promoter region of a downstream long TIF (as shown for gene GABPB1-AS1 in Supplementary Figure S14). Another type (30%) consists of two relatively longer TIFs that overlap at low degree. Polyadenylation cleavage sites of the upstream TIFs in both types typically occur within 2 Kb downstream of the next TIF start sites, which matches the observation in all tandem overlapping TIFs (Figure 3B). In order to examine potential regulatory crosstalk among overlapping transcripts, we measured pairwise gene co-expression pattern in 21 CML patients RNA-Seq data (26). As RNA polymerases usually goes further after poly(A) cleavage sites, we included the nonoverlapping transcripts in the upstream 10 Kb region. The TIF pairs are classified into different groups according to their overlap widths or the distance between two TIFs. Expression of TIFs are represented by its corresponding genes in the CML patients’ data. In general, the overlapping transcripts shows a slightly less negative correlation than non-overlapping pairs (Supplementary Figure S15). However, no significant difference in co-expression was detected among groups.

Having an advantage of capturing both transcription boundaries in overlapping transcripts, TIF-Seq2 is able to pinpoint interdependence of TSSs and PASs across overlapping transcripts (Supplementary Figure S14). In addition, we observed a type of TSS-PAS combination from tandem neighbouring genes, thus generating fused genes caused by transcriptional read-through.

### Linking transcription boundaries facilitates the identification of read-through transcripts

An extreme case of overlapping transcriptional units, is the case of those where we identified novel combination between TSS-PAS from neighbouring genes. As those events suggest transcriptional read-through and the potential generation of fused genes, we decided to study them in detail. Transcriptionally fused genes are in general challenging to identify by conventional RNA-Seq approaches (36–38). Gene fusions play a key role in oncogenesis (39), and although most known fusion transcripts arise through chromosomal rearrangements, they can also arise via transcription-induced chimeras (*e.g*. read-through or alternative splicing) (36). Short-read RNA-Seq can detect read-through transcripts by using the small proportion of reads connecting two neighbouring genes (37). However, this becomes extremely challenging when the fusion event involves an intermediate “stepping stone” exon not included in the annotations of the involved genes (*e.g*. LHX6-NDUFA8_03, Figure 3A). Additionally, RNA-Seq does not allow to identify which TSS of the upstream gene is connected to a particular PAS of the downstream gene. On the contrary, all TIF-Seq2 reads have the potential to detect such fusion events (not only the small fraction covering a splicing between the annotated genes) and define their complete boundaries (TSS to PAS). We discovered 29 non-annotated read-through transcripts in K562 cell lines, and once defined their boundaries, we were able to identify supporting splicing reads for all of them using RNA-Seq (25) (Supplementary Table S7). However, without the additional information provided by TIF-Seq2, those same RNA-Seq datasets were not sufficient to confidently classify them as read-through transcripts. Interestingly, during the preparation of this work, a few cases have been investigated in detail and independently reported as fusion transcripts in agreement with our findings (40, 41).

To investigate up to what degree alternative splicing contributes to the appearance of those read-through transcripts, we further validated 10 candidates using the TIF-Seq2 derived boundaries and amplicon-based enrichment of full-length isoforms followed by long-read sequencing (as before, Supplementary Figure S12). This revealed an interleaved and complex transcriptome organization (Figure 4A, B and Supplementary Figure S16).

**Figure 4.**
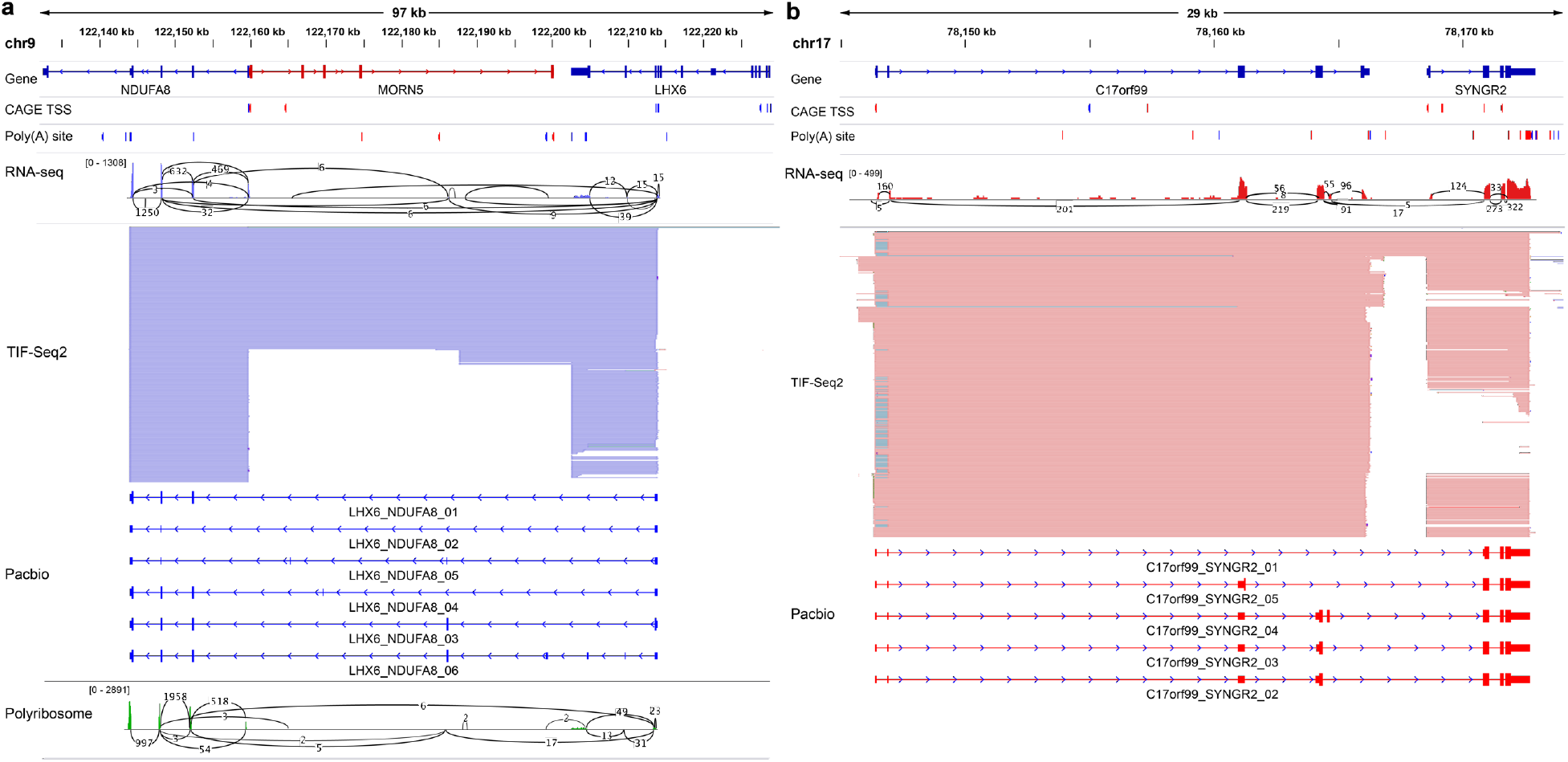
TIF-Seq2 facilitates the identification of read-through transcripts. (A) *LHX6-NDUFA8* read-through gene; (B) *C17orf99-SYNGR2* read-through gene. Annotated genes are listed on the top, followed by CAGE TSS and poly(A) site (PAS) tracks (12, 13). RNA-Seq validates the presence of splicing junctions (black lines with supporting number of reads) linking two adjacent genes. TIF-Seq2 track shows the transcriptional fusion events between adjacent genes (pink on the forward strand and purple on the reversed strand). PacBio Long-read sequencing of target transcripts validates the intergenic splicing events and dissect the transcription model of read-through genes. Polyribosome-associated RNA-Seq (27) data are labelled in green, with splicing junction in black lines between two genes, showing the coding potential of two transcript isoforms of *LHX6-NDUFA8*.

In some cases, read-through transcripts connect the 5’ regions of one gene with the coding sequence of the downstream one (e.g. C17orf99_SYNGR2_01, Figure 4B), suggesting a regulatory rewiring of the downstream gene (*e.g*. putative regulators of C17orf99 could regulate SYNGR2 expression), while in other cases, the read-through transcripts have the potential to encode fusion proteins (*e.g*. SPN_QPRT_04, Supplementary Figure S16). Our approach can detect even more complex scenarios, as is the case of transcripts initiating in the body of one gene but using the PAS of a downstream gene (e.g. *LHX6-NDUFA8* in Figure 4A). Or other complex cases where the read-through transcript connects both genes using a nonannotated exon (*e.g*. LHX6-NDUFA8_03-06 or C17orf99_SYNGR2_04 in Figure 4).

To provide additional supporting evidence for the identified read-through transcripts, we investigated their potential to encode fusion proteins. We used RNA-Seq from polyribosome associated mRNAs (27) and confirmed that reads connecting the novel splicing sites predicted by our long-read experiments can be identified in 10 read-through genes (Figure 4A). This suggests that the identified transcripts associate to polyribosomes, and thus have the potential to encode fusion proteins. In addition, we investigated their presence in CML patients by re-analysing clinical RNA-Seq data (26). Even if we had to restrict our analysis to those few reads bridging the gene pairs, we were able to confirm their expression in patients (Supplementary Figure S17). Thus, by combining an improved annotation using TIF-Seq2 with available clinical short read dataset, we were able to identify read-through transcripts that would be ideal candidates for in deep molecular characterization in disease models.

## Discussion and conclusion

Here we have presented an improved approach, TIF-Seq2, designed to link transcription boundaries in complex genomes. This approach is in agreement with previous maps of TSS and PAS, and can enrich current transcriptome studies by clarifying complex arrangement of overlapping transcripts. By focusing its sequencing power on TSS and PAS, it can easily define the complete boundaries of lncRNAs or other lowly expressed transcriptional features. This allows also to link genes with putative regulatory regions and to facilitate its genetic manipulations (*e.g*. using CRISPRi or CRISPRa). As TIF-Seq2 is based in shortread sequencing, in principle it could be easily combined with traditional target enrichment approaches to link unannotated, but experimentally validated, TSS or PAS sites (*i.e*. using a unique probe targeting TSS or PAS of interest, without the need to enrich the intervening region). Therefore, it opens the door for the design of target enrichment strategies to allow their study with other sequencing approaches. By combining TIF-Seq2 information with RNA-seq, we show how an improved transcriptome annotation can refine our analysis of available clinical RNA-Seq datasets. This would be especially valuable to prioritize the transcriptional features more relevant for in depth molecular validation. Application of TIF-Seq2 goes beyond the study of the human genome, and it could be particularly useful to facilitate the annotation of less studied genomes.

TIF-Seq2 is particularly useful to dissect those overlapping transcripts which pose a challenge to investigate by any other short-read RNA-Seq approaches. It can easily distinguish overlapping transcript units from each other and quantify their expression, thus facilitating the interpretation of isoform-specific response in different conditions. By examining the interaction between overlapping transcripts, we show that partially overlapping transcripts often use poly(A) sites within 2 Kb of the TSS of the downstream genes. This suggests that during this window the cellular machinery is able to distinguish between short transcripts that should be terminated from those that will proceed to produce a full RNA. This is reminiscent of the mechanism previously described for PROMPTs and the transcription between closely spaced promoters (20, 42). We show that partially overlapping transcripts are formed by two main classes of upstream transcripts: those originating proximal to the downstream TSS (less than 1Kb) and a second-class involving transcript originating upstream. Those arrangements are difficult to study by RNA-Seq and CAGE and easily lead to the misassignment of reads corresponding to the upstream promoter to the downstream transcription unit. Finally, we identified read-through transcripts by linking the usage of particular TSS and PAS from neighbouring genes. This reveals the ability of read-through transcripts to putatively rewire their regulation (*i.e*. usage of alternative shared upstream promoters) and produce fusion proteins. We confirmed the identified fusion transcripts by targeted long-read sequencing and confirmed their putative coding potential by reanalysing ribosome-profiling data. We show that the identified transcripts can also be observed in clinical RNA-Seq datasets. Our work also points to the intrinsic limitation of RNA-Seq, which in some cases needs to be complemented by alternative approaches to resolve complex overlapping transcriptional structures. In conclusion, we think that TIF-Seq2 has great potential to complement the current transcriptomic approaches, help dissect the overlapping transcriptome and thus fill in missing puzzle pieces to improve our understanding of transcription regulation.

## Supplementary Data

Figure S1. Detailed TIF-Seq2 workflow.

Figure S2. Detail of simultaneous sequencing of 5’and 3’reads.

Figure S3. Viability plot for K562 treated with Imatinib.

Figure S4. TIF-Seq2 data analysis process.

Figure S5. Size distribution of TSS and PAS clusters width.

Figure S6. Distribution of poly(A) site distance between TIF-Seq2 and 3’T-filling sequencing.

Figure S7. Distance between TIFs and Gencode transcripts.

Figure S8. Distribution of TIF counts per annotated genes.

Figure S9. Category of TIFs.

Figure S10. Differential expression of TIFs before and after Imatinib treatment.

Figure S11. Differential expression comparison between TIF-Seq2 and 3’T-fill.

Figure S12. Design of Pac Bio long-read target sequencing.

Figure S13. An unannotated transcribed region identified by TIF-Seq2.

Figure S14. Interdependency of TSS and PAS in GABPB1-AS1.

Figure S15. Measurement of gene co-expression based on the overlapping pattern of tandem TIFs.

Figure S16. Example of a transcriptionally fused molecules connecting SPN and QPRT.

Figure S17. Splicing junctions across read-through genes can be detected in in CML patients.

Table S1. List of oligos used for TIF-Seq2 library preparation and target enrichment.

Table S2. Summary of read pairs in TIF-Seq2.

Table S3. Read alignment and library complexity.

Table S4. Non-overlapping unannotated transcribed regions.

Table S5. Transcription isoforms in K562 cells identified by TIF-Seq2.

Table S6. All poly(A) sites identified by 3’T-fill sequencing and differential expression measurement using DESeq2.

Table S7. Splicing junctions linking read-through transcripts in an independent K562 cells RNA-Seq data (GSE105161).

Table S8. New transcription features validated by long-read sequencing in BED format.

## Data availability

All TIF-Seq2, 3’T-fill sequencing and long-read sequencing files were deposited in the Gene Expression Omnibus under accession number (GSE140912).

The source code for TIF-Seq2 data analysis is available on GitHub (https://github.com/jingwen/TIFseq2)

## Acknowledgements

We thank Donal Barrett for technical assistance, Yuanyuan Xi for supporting computational analysis, and all members of Pelechano, Kutter and Friedländer laboratories for discussion. We thank Aino Järvelin and Judith Zaugg for early discussion. We would like to thank Dr. Andreas Schreiber for his help with the access to clinical RNA-Seq datasets. The authors would like to acknowledge support from Science for Life Laboratory, the National Genomics Infrastructure (NGI) and Uppmax for providing assistance in massive parallel sequencing and computational infrastructure.

## Author Contributions

VP, BL, JW, WW and LS conceived the study. BL and SM performed experiments and assisted with data interpretation. JW, BL, WW and VP designed data analysis and interpreted data. JW performed computational analysis. JW, BL and VP drafted the original manuscript. All authors reviewed and edited the manuscript. VP supervised the whole project.

