## Supplementary Figures for "Dissecting chronic myeloid leukaemia overlapping transcriptome with TIF-Seq2"

### Supplementary Figures and Legends

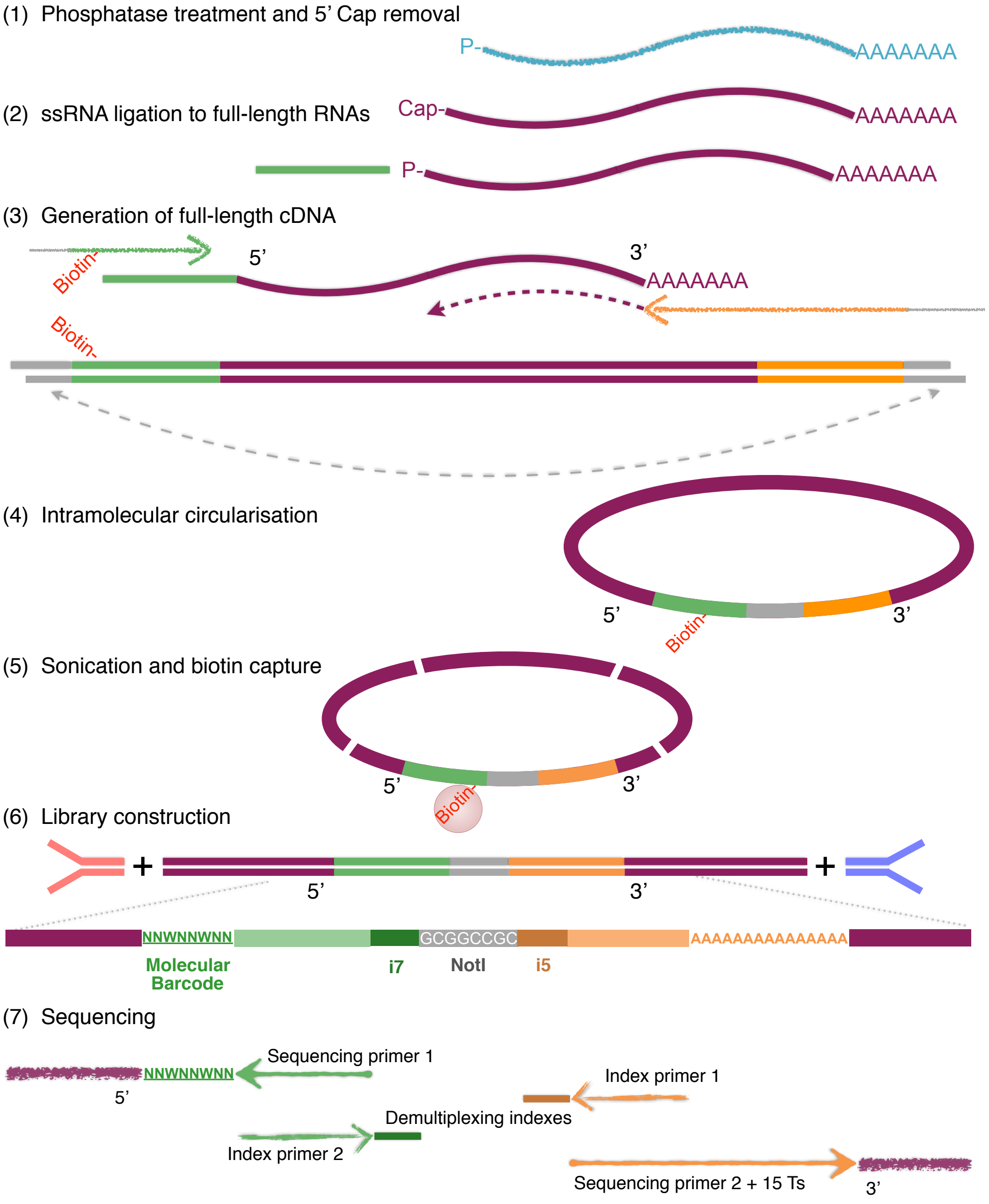


**Supplementary Figure 1. Detailed TIF-Seq2 workflow.** (1)Total RNA was dephosphorylated by incubating with Alkaline Phosphatase, Calf Intestinal(CIP). mRNAs were decapped by incubating with Cap-Clip which exposed the 5' phosphomonoester of mRNA and allowed for ligation to RNA adaptor. (2) RNA was ligated overnight with DNA/RNA chimeric oligonucleotide adaptor using T4 RNA ligase. (3) mRNAs were reverse transcribed using barcoded oligo-dT primers. First strand cDNA was used as the template for PCR amplification. PCR products from different samples were pooled and subjected to NOT1-HF endonuclease digestion. (4) To favour intramolecular ligation, digested PCR products were highly diluted and ligated with a high concentration of T4 DNA ligase. plasmid-safe was used to remove the unligated linear PRC products. (5) Circularised cDNAs were fragmented via focused acoustics. Biotin labelled fragments which contained the 5' and 3' connecting site was enriched using Dynabeads M280 streptavidin. (6) Following end repair and dA tailing, each sample was ligated to TIF-Seq2 annealed adaptors. Samples were amplified, size selected and sequenced with the depicted oligos with expected size distribution in the range of 300bp-1000bp. Purified DNA library loaded onto NextSeq 500 sequencer with custom primers and custom settings (read1 76bp, read2 76bp, index1 6bp and index2 6bp).

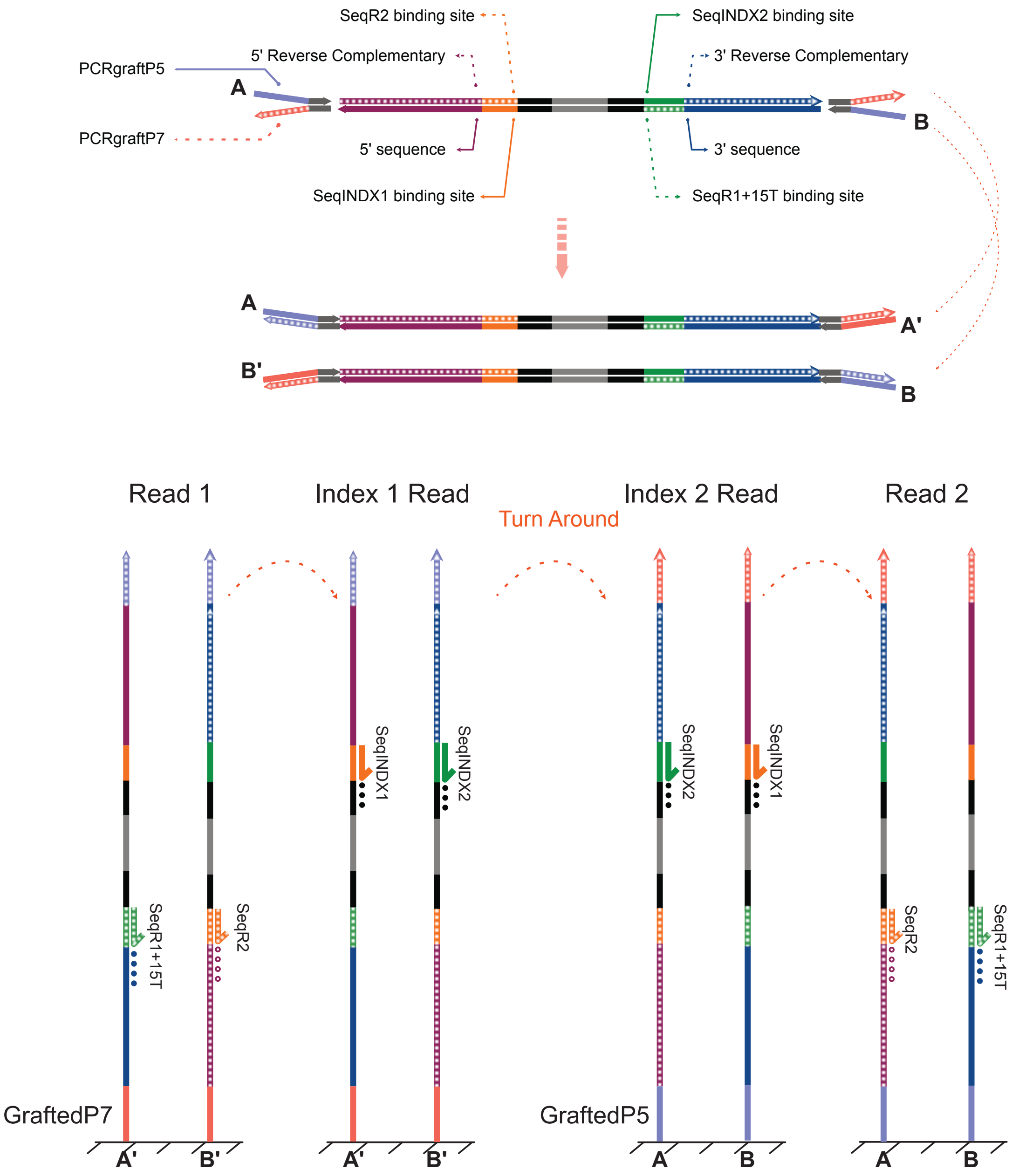


**Supplementary Figure 2. Detail of simultaneous sequencing of 5´and 3´reads.** a, Compared to standard Illumina reads structure, in TIF-Seq2 reads the flow-cell grafting region is decoupled from the sequencing primer regions. This will generate in the flow cells clusters with the same graft sequences but reverse complementary internal part. After cluster generation only one strand is kept attached to the flow cell to allow sequencing by synthesis. In TIF-Seq2, half of the clusters will contain single-stranded DNA reverse complementary to the first sequencing index (A) and the other half reverse complementary to the second sequencing index (B). By adding simultaneous both sequencing indexes both clusters A and B are sequenced at the same time. As we used different sequencing index for the 5´ and 3´ regions, we can correctly classify TSS and PAS during the analysis. b, detail of the sequencing cycles in the Illumina flow-cell. Please note that in some cases the same sequence (and not the reverse complementary) is present, preventing thus the annealing of the sequencing oligos.

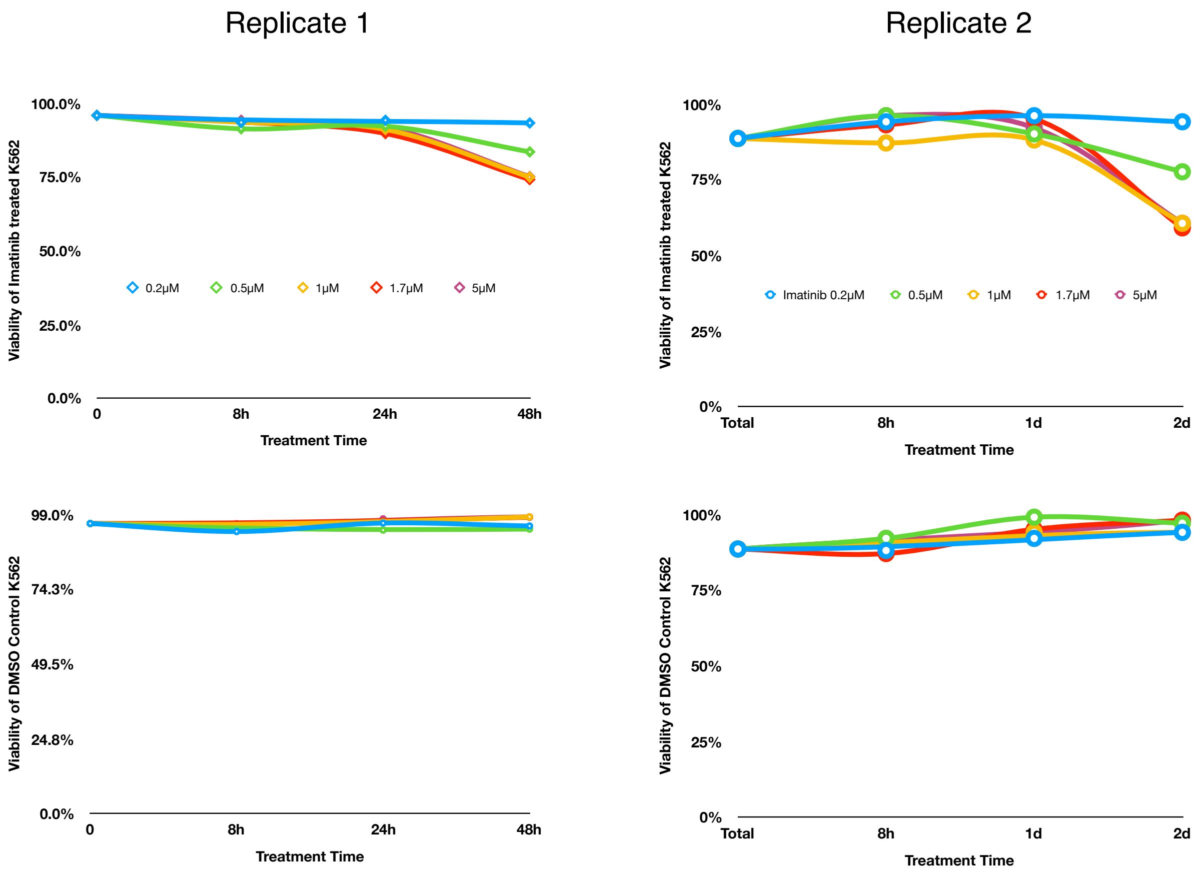


**Supplementary Figure 3. Viability plot for K562 treated with Imatinib.** a, Two biological replicates of K562 cells treated with series of concentrations (0.2 0.5, 1, 1.7, 5 $\mu$M) Imatinib for 8, 24 and 48 hours. Viability was estimated by EVE automated cell counter. b, matched controls corresponding DMSO control treatment. We selected 1 $\mu$M Imatinib treatment for 24 hours as it causes a clear phenotype at 48h hours, but the viability is still high at 24h.

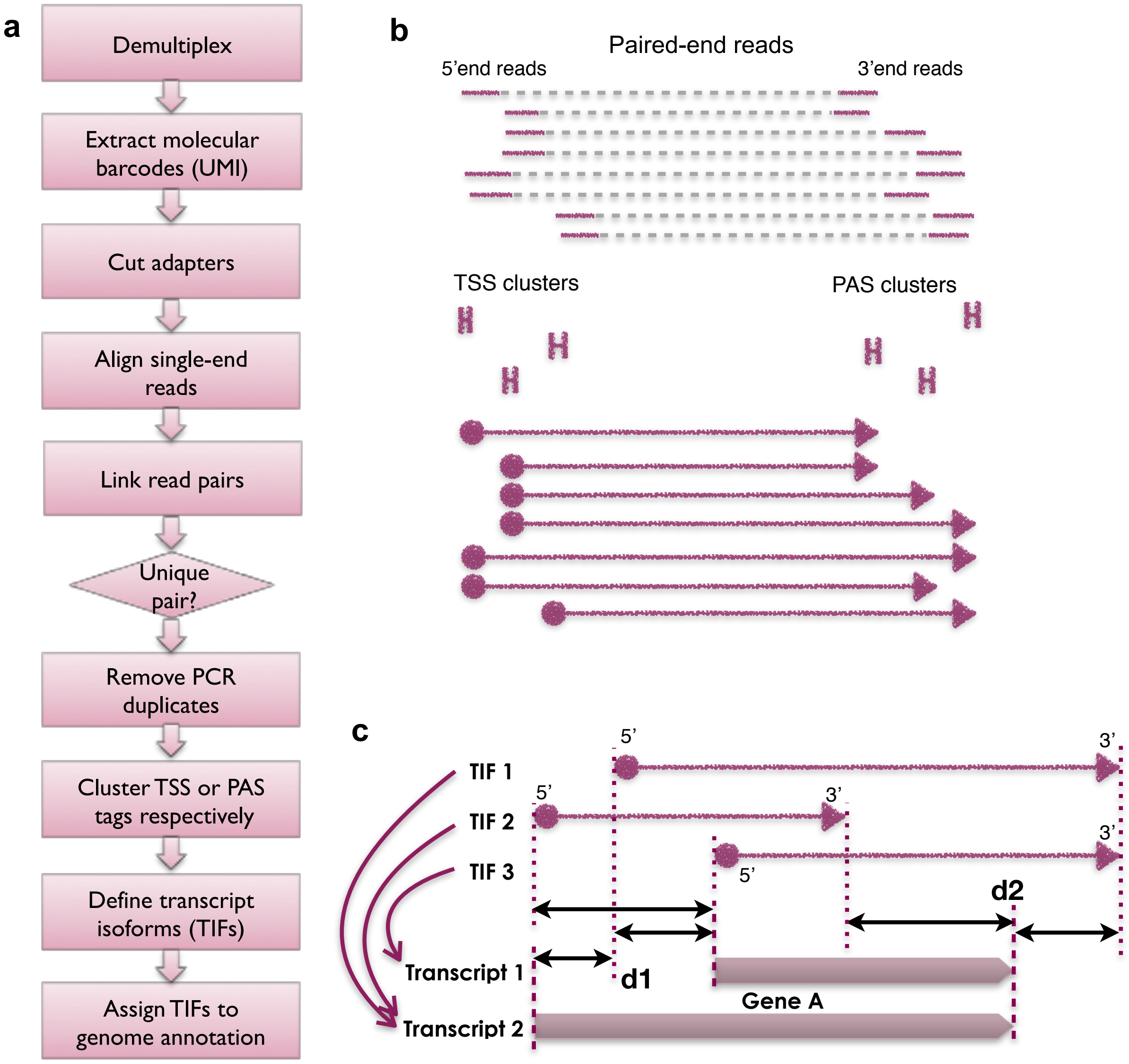


**Supplementary Figure 4. TIF-Seq2 data analysis process.** (A) Data analysis workflow of TIF-Seq2. (B) Transcription start site (TSS) tags and poly(A) site (PAS) tags were clustered independently. Two tags located within 10 nt were cluster together. After forming clusters at the transcription boundaries, we linked the TSS and PAS clusters according to TIF-Seq2 read pairs. (C) Assignment of TIFs to annotated transcripts and genes. TSS distances (d1) and PAS distances (d2) are calculated between a TIF and its overlapping annotated transcripts. The TIF is assigned to the transcript with the least sum of d1 and d2 among all overlapping transcripts, further assigned to the gene that harbour the transcript.

A

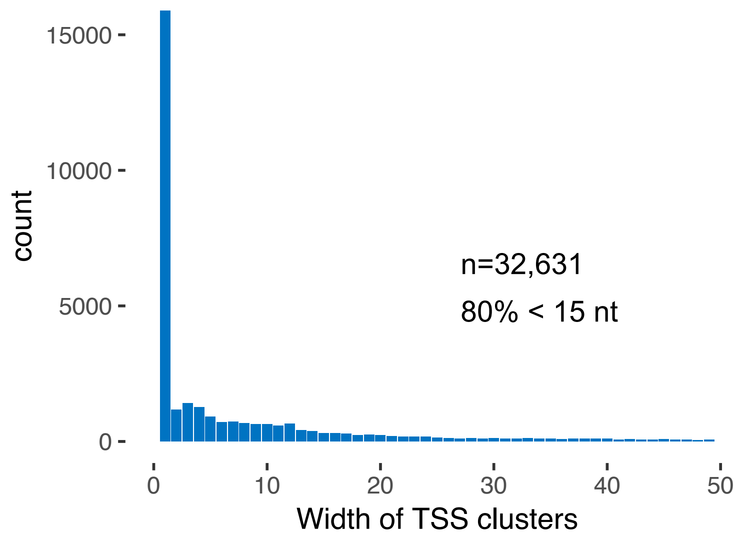


B

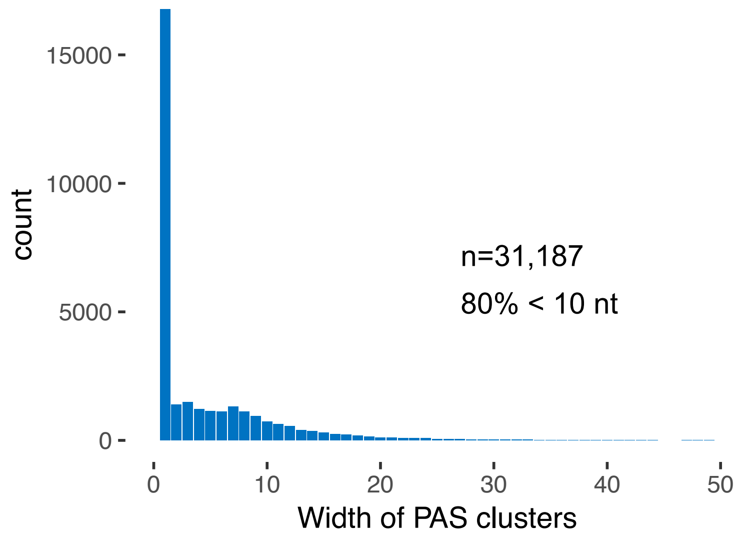


**Supplementary Figure 5. Size distribution of TSS and PAS clusters width.** (A-B) We identified 32,361 clusters of TSS (A) and 31,187 clusters of PAS (B) with TIF-Seq2 in K562 cells. The distributions of their widths show that 80% of TSS clusters are shorter than 15 nt and 80% of PAS clusters are shorter than 10 nt.

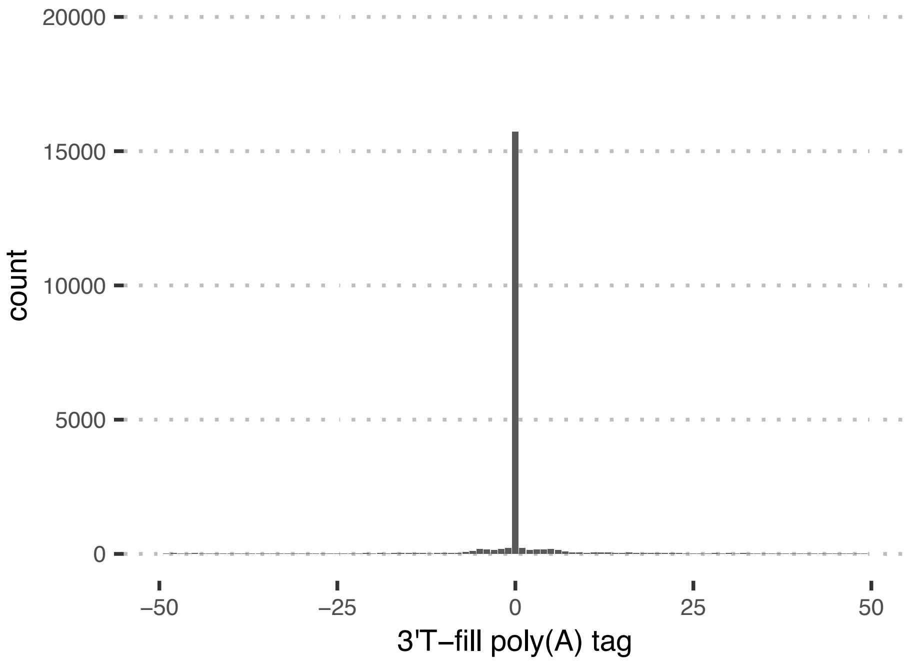


**Supplementary Figure 6. Distribution of poly(A) site distance between TIF-Seq2 and 3'T-filling sequencing.** Poly(A) sites determined by TIF-Seq2 agree with poly(A) tags determined by 3'T-fill sequencing. The distances are calculated from the peaks of 3'-end clusters in TIF-Seq2 to their nearest 3'-end tags in 3'T-fill sequencing.

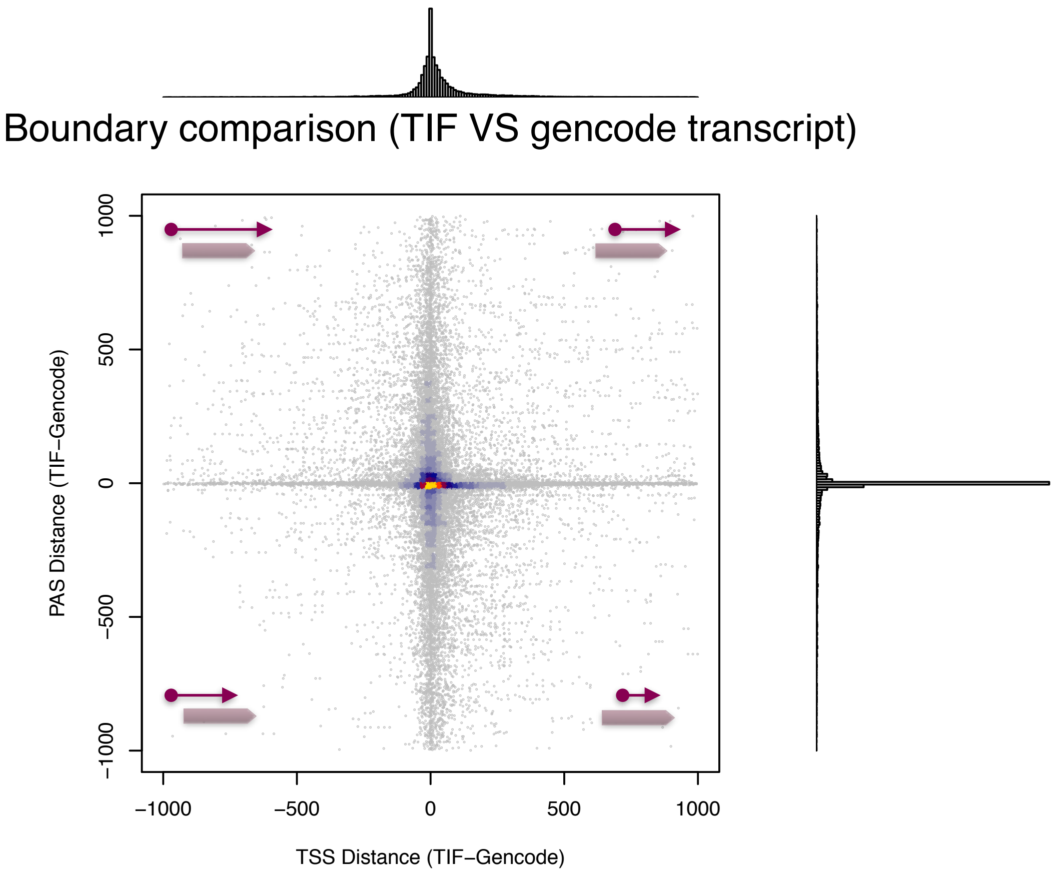


**Supplementary Figure 7. Distance between TIFs and Gencode transcripts.** Comparison of TIF boundaries and annotated transcript boundaries (Gencode v28). TIFs were assigned to the annotated transcripts according to Supplementary Figure 4c. TSS distance (x axis) and PAS distance (y axis) demonstrate that a large number of TIF-Seq2 defined transcript boundaries cover the entire annotated transcripts.

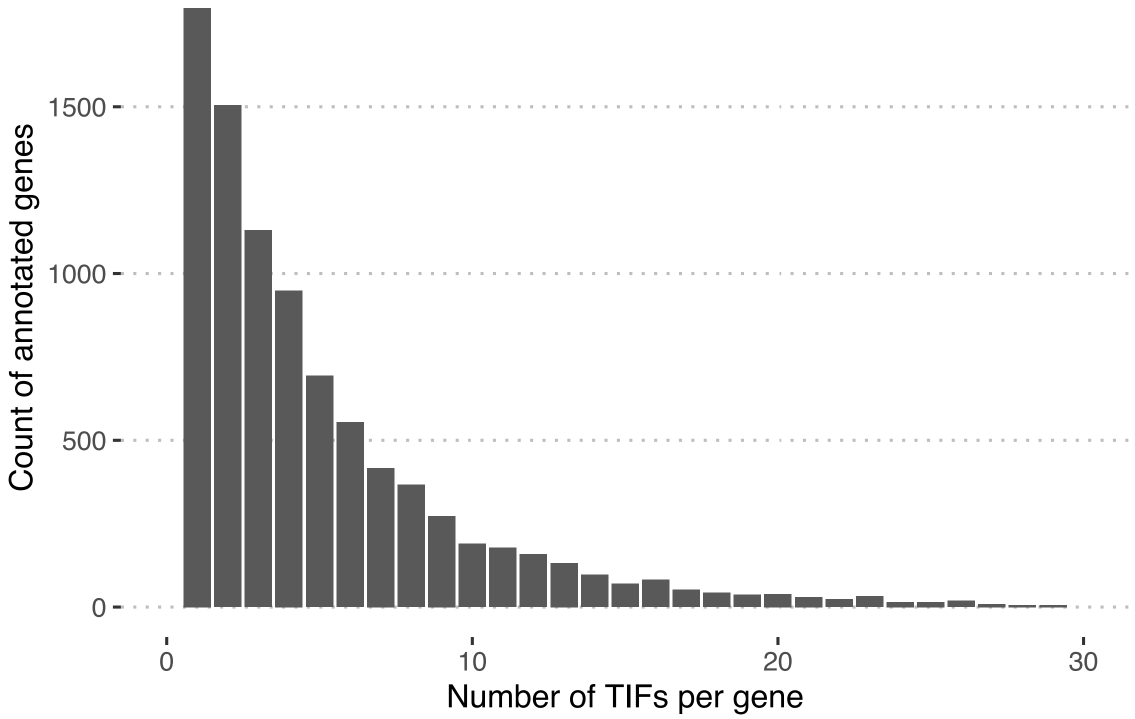


**Supplementary Figure 8. Distribution of TIF counts per annotated genes.** TIFs were assigned to the annotated genes (Gencode v28) according to Supplementary Figure 4c. Among 9006 Gencode genes which is covered by TIF-Seq2, 7210 genes have more than one transcript isoform boundaries (TIFs). With current depth of our sequencing run, there are on average 4 TIFs per gene.

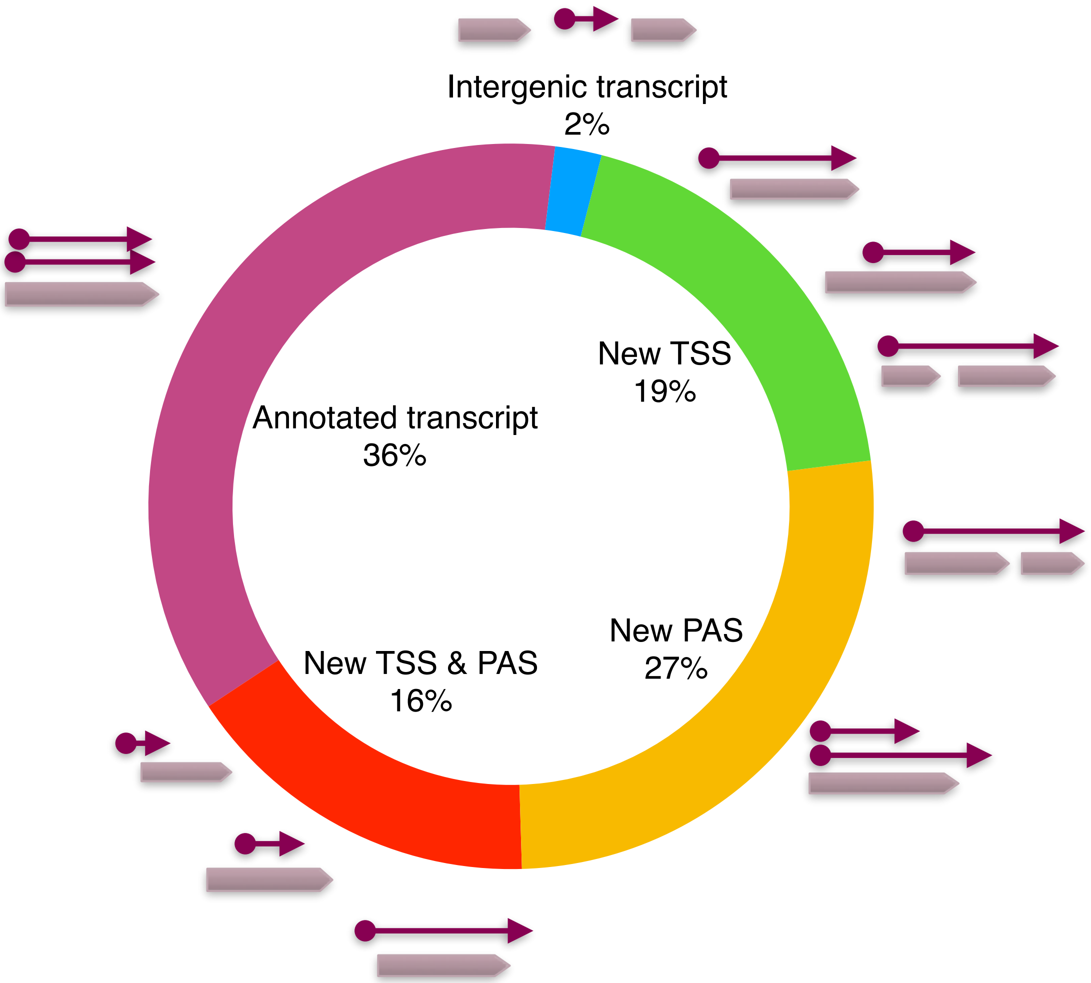


**Supplementary Figure 9. Category of TIFs.** Transcription isoforms (TIFs) are assigned to the human genome annotation Gencode v28. If both transcription boundaries in a TIF are located within 200 bp of annotated transcripts boundaries, the TIF is assigned to annotated transcript. Otherwise, it is a new transcript, although it may overlap with an annotated gene. The TIFs that don't overlap with any genes are classified as intergenic transcript.

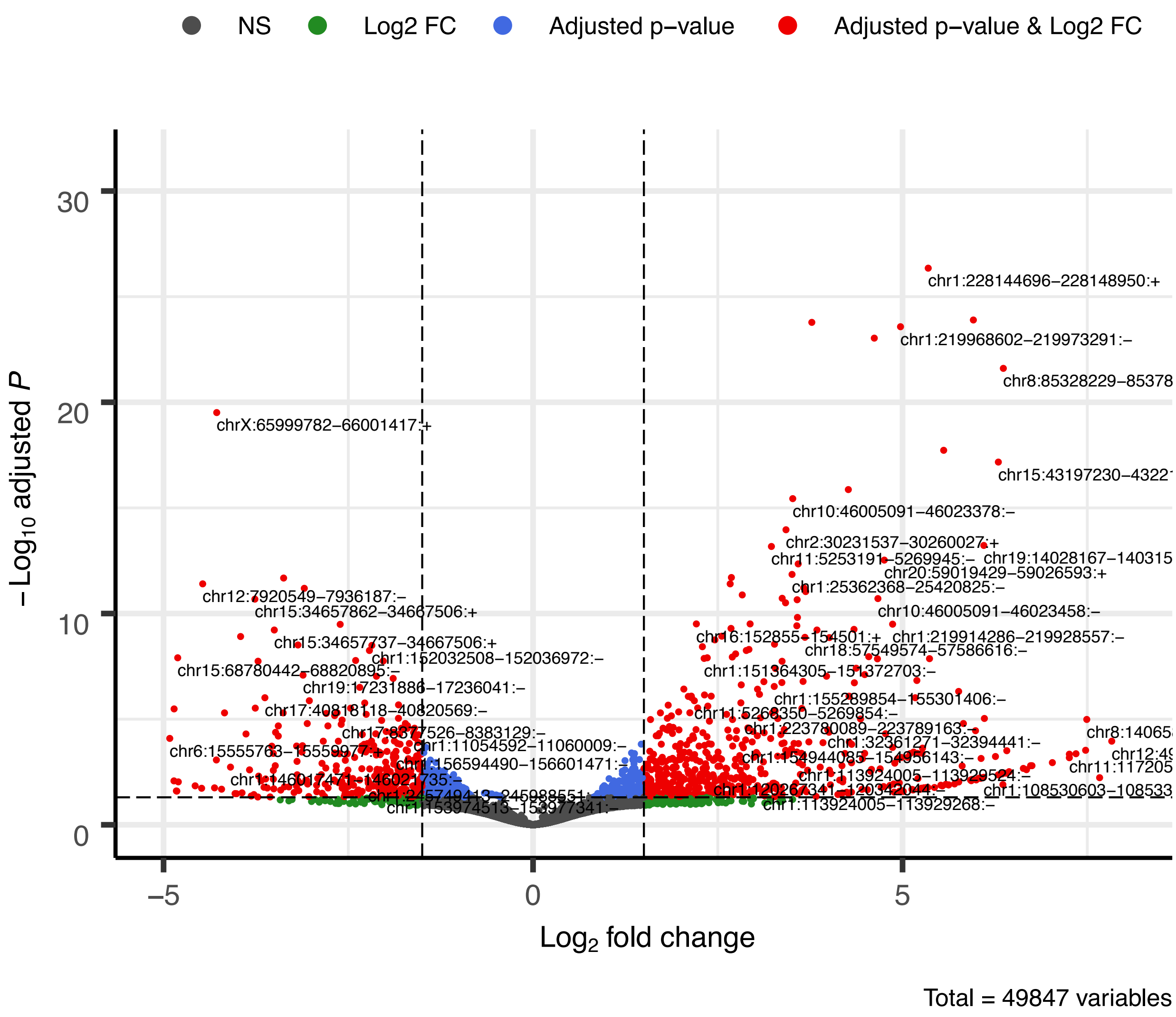


**Supplementary Figure 10. Differential expression of TIFs before and after Imatinib treatment.** We identified 49,847 transcript isoforms (TIFs) by using TIF-Seq2. Each dot represents differential expression of one TIF before and after Imatinib treatment. The threshold of significant level is $Log_2$-scale fold change over 1.5 or below -1.5, and adjusted p-value less than 0.05

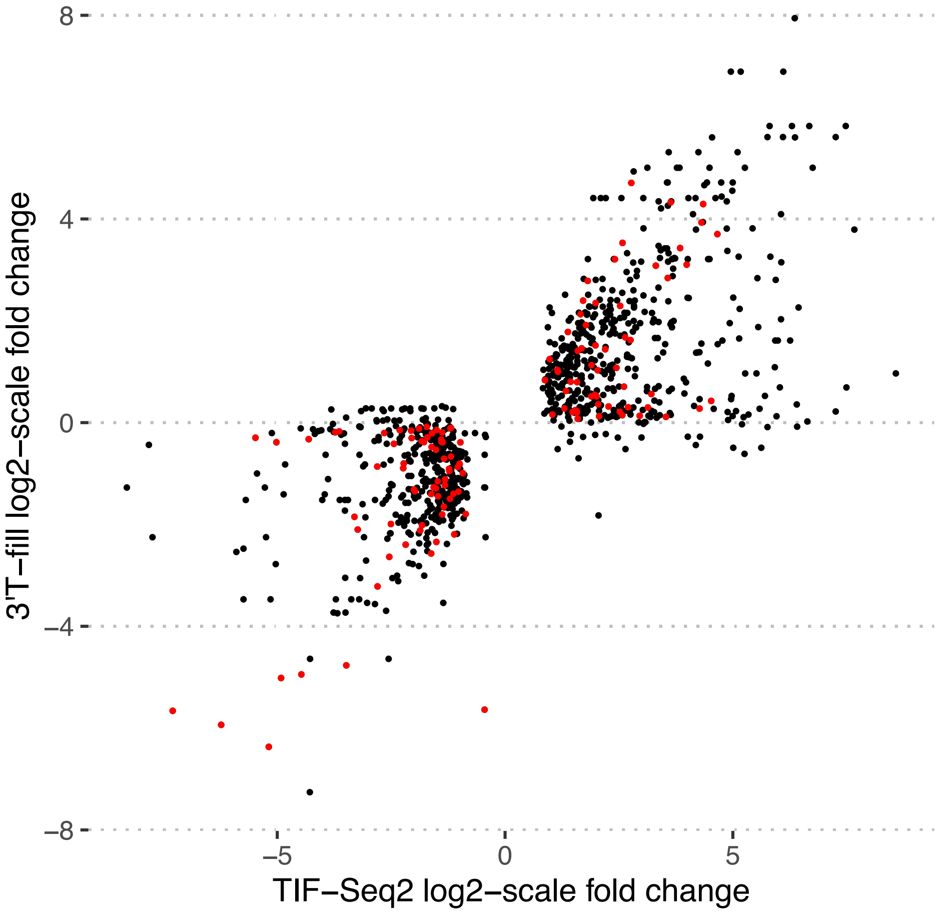


**Supplementary Figure 11. Differential expression comparison between TIF-Seq2 and 3´T-fill.** Comparison of $log_2$-scale fold changes between differential expressed (adjusted p-value <= 0.05) TIFs (x-axis) and their corresponding 3'end tags measured by 3'T-fill sequencing (x-axis). As one PAS can correspond to multiple identified TIFs, we indicate those 3' end tags assigned unequivocally to an individual TIF in red. The black dots represent 3' end tags shared by more than one TIF-Seq2 identified isoform. That explains why some 3' end tags apparently not differentially expressed when considering 3´T-fill, can be identified as regulated when considering the matched TSS.

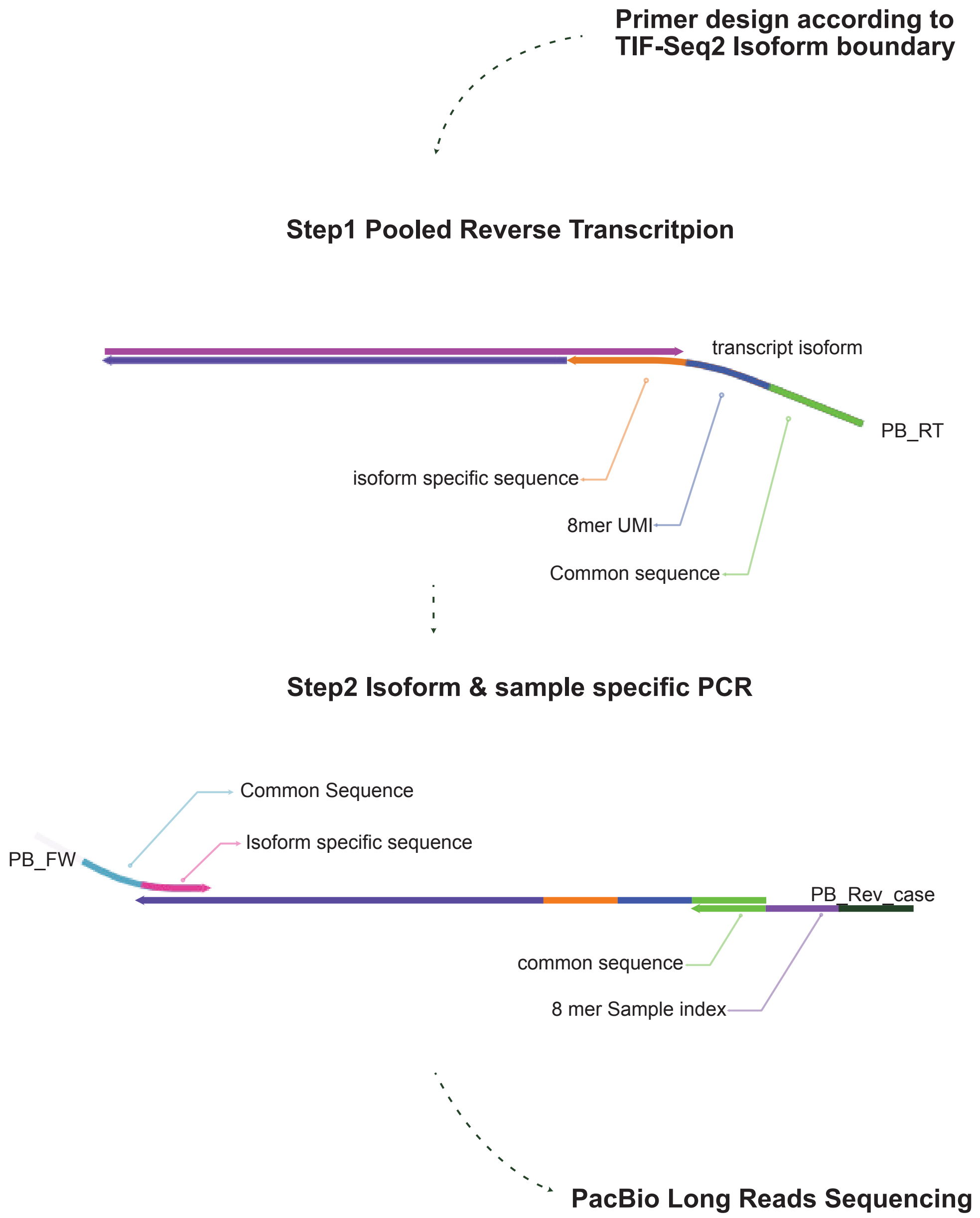


**Supplementary Figure 12. Design of Pac Bio long-read target sequencing.** We used TIF-Seq2 derived information to designed the required primers for target amplification (see methods and Supplementary Table S6). Target mRNAs were reverse transcribed using a pool of Pooled reverse transcription. We pooled equal amount of all isoform-specific primers (PB_RT) and used total RNA as a template for reverse transcription. cDNA was split in individual wells and subjected to PCR targeted amplification using PB_FW and PB_Rev oligo. PCR products were purified, pooled and used for Pac Bio sequencing.

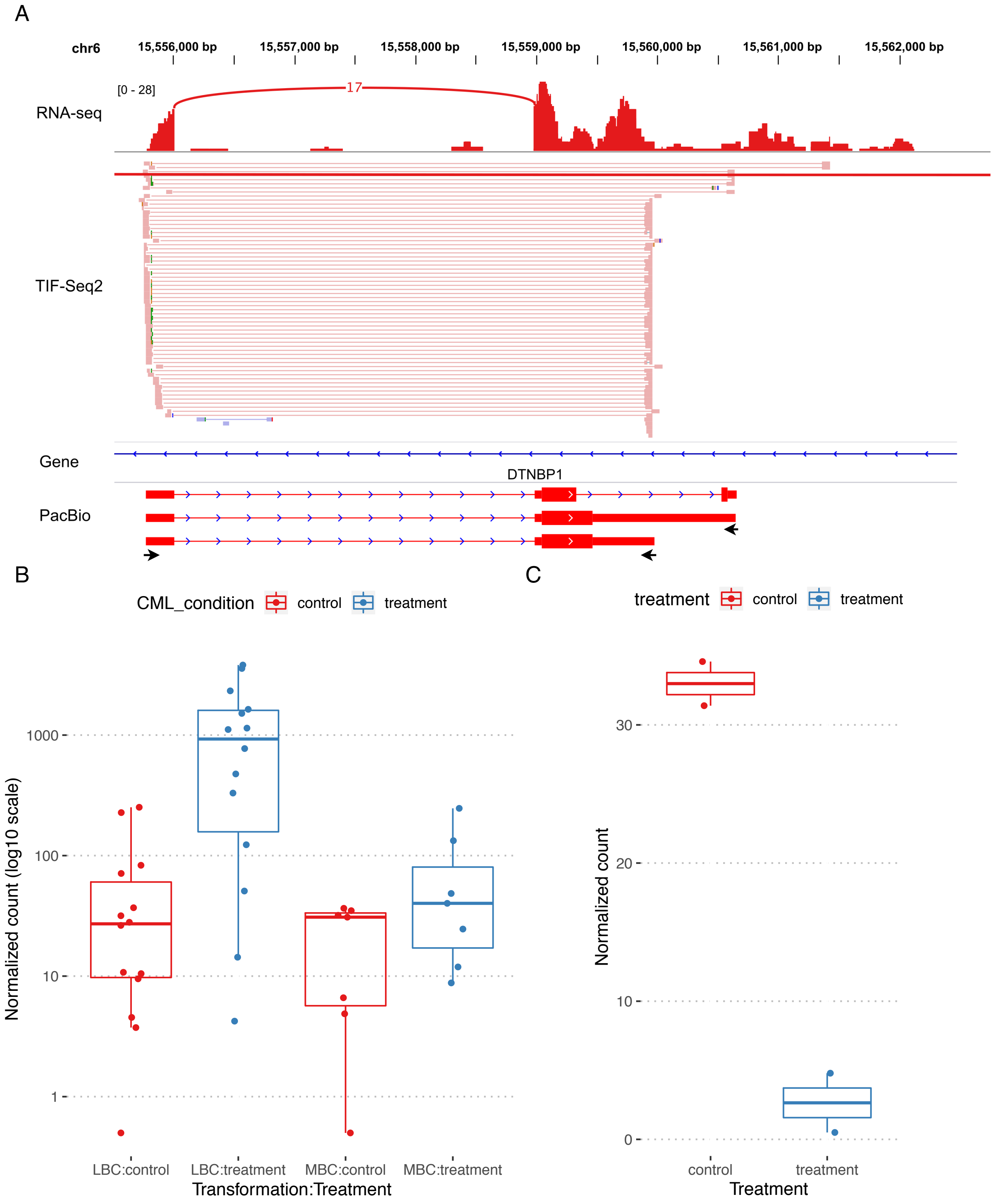


**Supplementary Figure 13. An unannotated transcribed region identified by TIF-Seq2.** (A) TIF-seq2 track (in pink) demonstrates that an unannotated transcribed region on chr6 has least three TIFs, supported by RNA-seq (on top). We designed primer pairs to enrich two target regions (marked by arrows) and sequence the whole amplicon by using PacBio long-read sequencing. (B) and (C), Gene expression before and after drug treatment *in vitro* and *in vivo*. In chronic myeloid leukaemia (CML) patients (B) who developed blast crisis (LBC or MBC), the unannotated gene was significantly upregulated ($log_2$ fold change = 3.67, adjusted p-value = 7.90 x $10^{-16}$) after Imatinib treatment, while in K562 cells (C) it was downregulated ($log_2$ fold change = -4.00, adjusted p-value = 7.94 x $10^{-4}$).

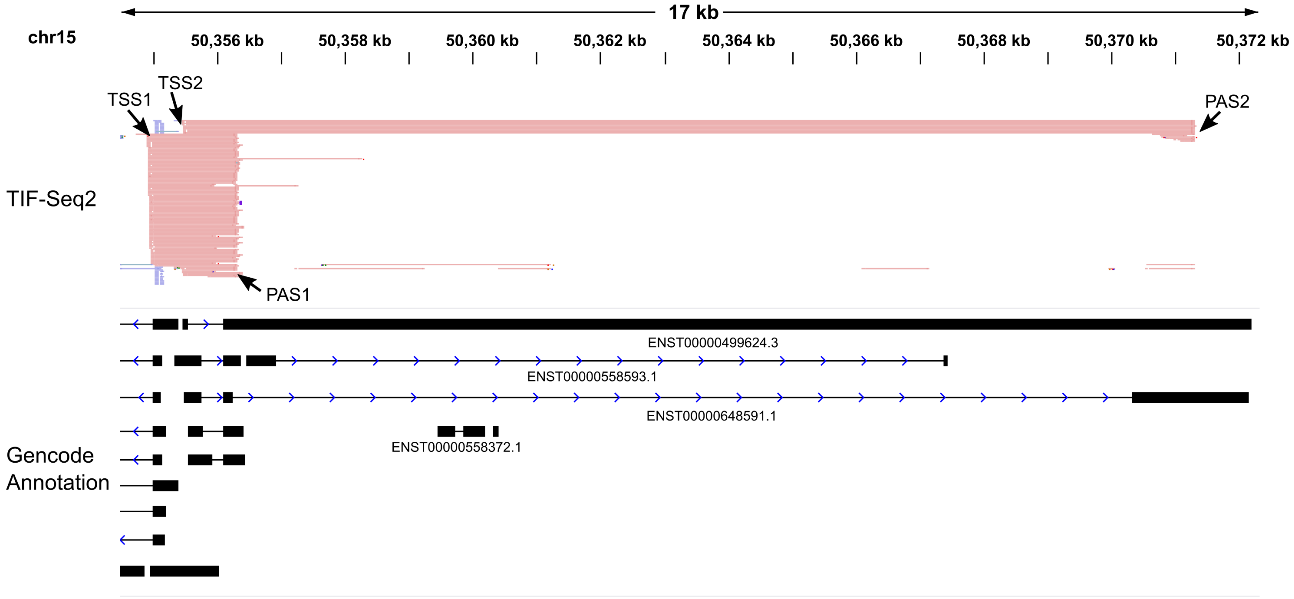


**Supplementary Figure 14. Interdependency of TSS and PAS in GABPB1-AS1**. Pink lines represent identified transcripts by TIF-Seq2. The overlapping transcripts on chr15 has two transcription start sites (TSSs) and two poly(A) sites (PASs). As TIF-Seq2 captures the co-occurrence of TSS and PAS, it is able to identify the interdependence of the transcription boundaries (TSS1-PAS1 and TSS2-PAS2), while the standard RNA-seq cannot.

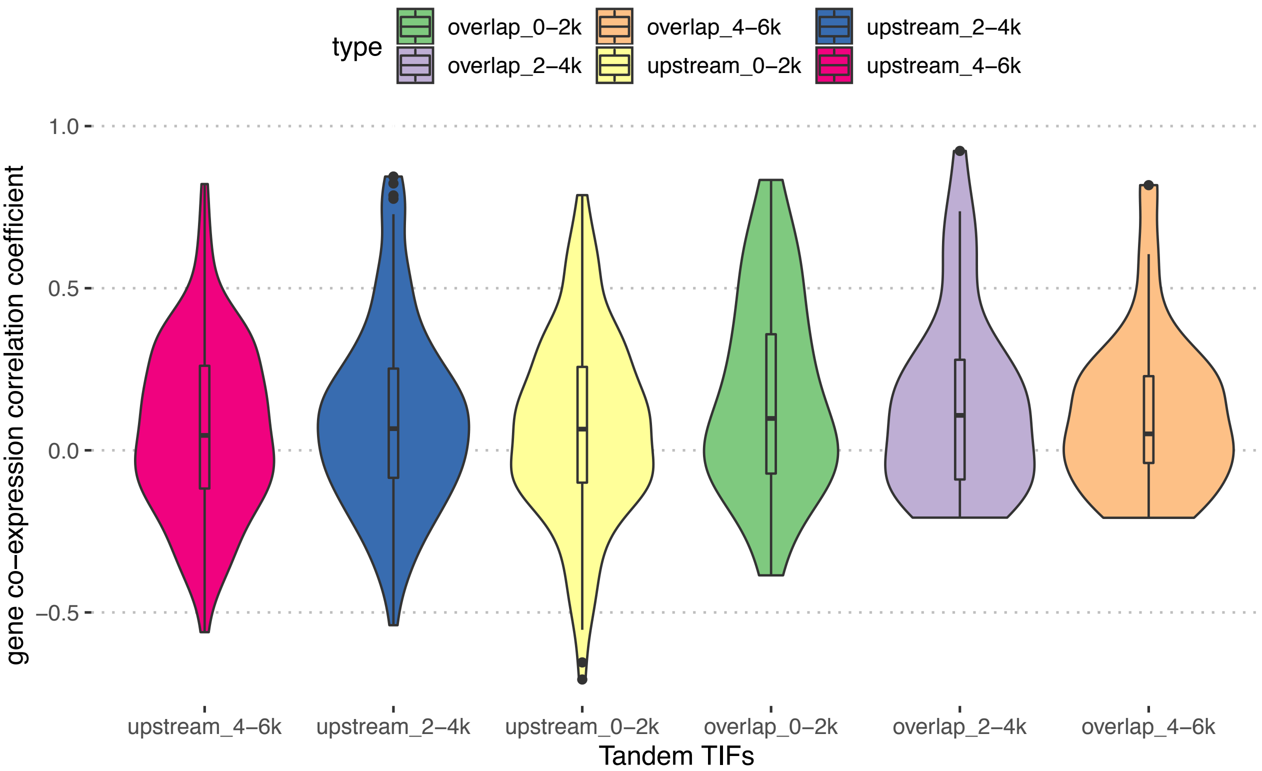


**Supplementary Figure S15. Measurement of gene co-expression based on the overlapping pattern of tandem TIFs**. The TIF pairs are classified into different groups according to their overlap widths (overlapping TIFs) or distances between two TIFs (non-overlapping TIFs). Expression of TIFs are represented by its corresponding genes in CML patients' RNA-seq data. Spearman correlation coefficient were calculated for each gene pairs.

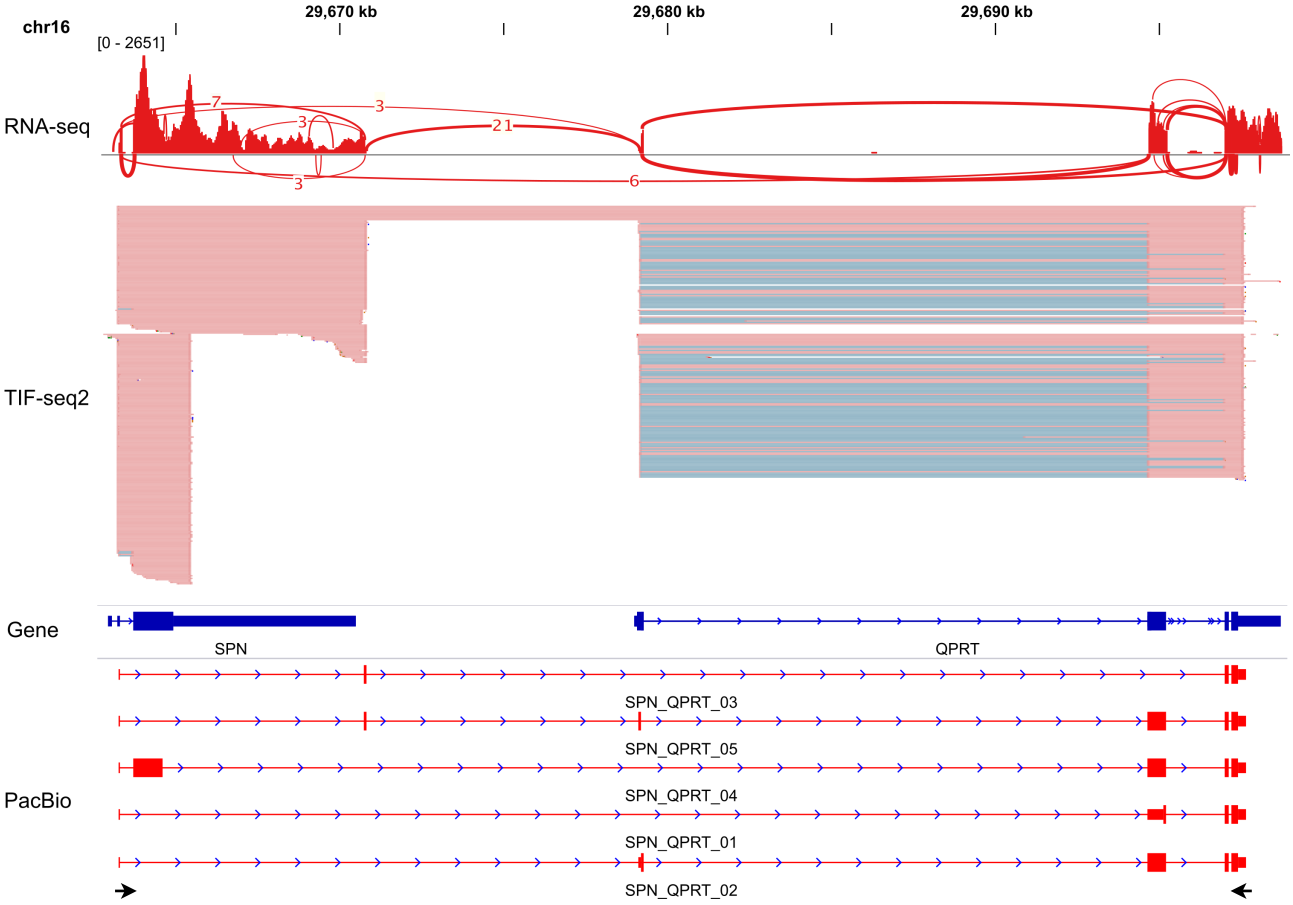


**Supplementary Figure 16. Example of a transcriptionally fused molecules connecting SPN and QPRT.** TIF-Seq2 (pink line) shows the transcriptional fusion events between SPN and QPRT. RNA-seq validates the presence of splicing junctions (red lines with supporting number of reads) linking them,. PacBio Long-read sequencing of target transcripts validates the intergenic splicing events and dissect the transcription model of SPN-QPRT read-through gene. Primer designed for target validation are marked in arrows.

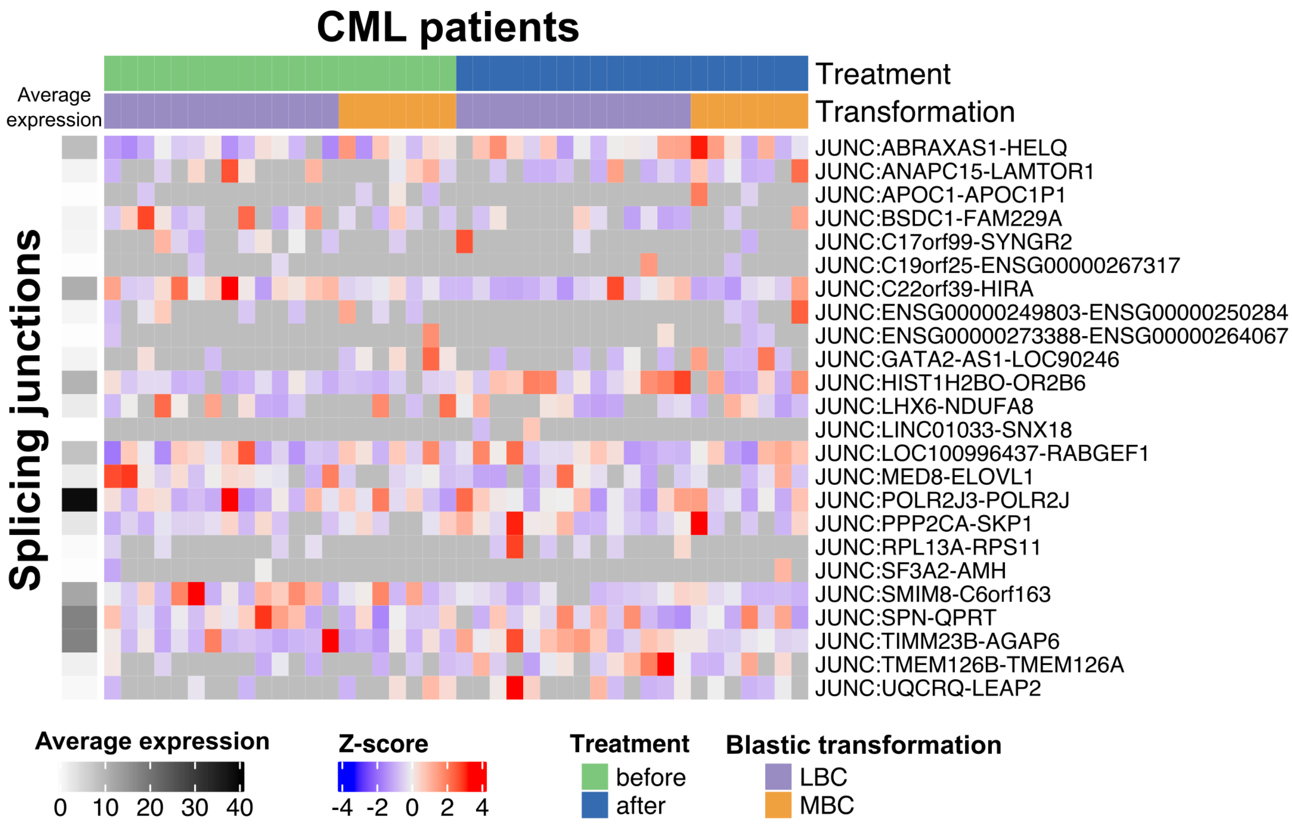


**Supplementary Figure 17. Splicing junctions across read-through genes can be detected in in CML patients.** The expression value was calculated as read counts containing splicing junctions that link two adjacent genes, quantified using featureCounts and normalized by DEseq2. Each row represents splicing junctions between adjacent genes, and each column is a patient before or after drug treatment. Each cell represents the Z-score of expression in each sample. No detectable expression is marked in grey. The left panel shows the average expression of the splicing junctions across all patient samples.

**Supplementary Table S1**. Oligos used for TIF-Seq2 library preparation and target enrichment
New transcription features validated by long-read sequencing in BED format. The predicted coding regions are listed in the 7th and 8th columns.

**Supplementary Table S2**. Numbers of read pairs from TIF-Seq2 run on K562 cells before and after Imatinib treatment

**Supplementary Table S3**. Number of aligned TIF-Seq2 read pairs and the read pairs left after removing PCR duplicates.

**Supplementary Table S4**. Non-overlapping unannotated transcribed regions. The 6th column shows the number of transcription isoforms (TIFs) identified by TIF-Seq2. The 7-10th columns show the normalised read counts in an independent RNA-seq study of K562 cells (GSE105161). The 11-16th columns are differential expression measurement by DESeq2.

**Supplementary Table S5**. Transcription isoforms (TIFs) in K562 cells identified by TIF-Seq2. The 5th column are the annotated genes which TIFs are assigned to. The 6-10th are the raw read-pair counts of each TIF. The last two columns are the differential expression measurement by DESeq2

**Supplementary Table S6**. All poly(A) sites identified by 3'T-fill sequencing and differential expression measurement using DESeq2 (including average expression, log2-scale fold change and adjusted p-value)

**Supplementary Table S7**. Splicing junctions linking read-through transcripts in an independent K562 cells RNA-seq data (GSE105161). The 5-9th columns show the raw read counts covering splicing junctions.

**Supplementary Table S8**. New transcription features validated by long-read sequencing in BED format.
